# Contribution of resident and circulating precursors to tumor-infiltrating CD8^+^ T cell populations in non-small cell lung cancer patients

**DOI:** 10.1101/2020.08.14.249789

**Authors:** Paul Gueguen, Christina Metoikidou, Thomas Dupic, Myriam Lawand, Christel Goudot, Sylvain Baulande, Sonia Lameiras, Olivier Lantz, Nicolas Girard, Agathe Seguin-Givelet, Marine Lefevre, Thierry Mora, Aleksandra M. Walczak, Joshua J Waterfall, Sebastian Amigorena

## Abstract

Tumor-infiltrating lymphocytes (TILs) in general, and CD8^+^ TILs in particular, represent a favorable prognostic factor in non-small cell lung cancer (NSCLC). The tissue origin, regenerative capacities, and differentiation pathways of TIL subpopulations, however, remain poorly understood. Using a combination of single cell RNA and T cell receptor (TCR) sequencing, we investigate the functional organization of TIL populations in primary NSCLC. We identify two CD8^+^ TIL subpopulations expressing memory-like gene modules: one is also present in blood (circulating precursors) and the other one in juxta-tumor tissue (tissue resident precursors). In tumors, these two precursor populations converge through a unique transitional state into terminally differentiated cells, often referred to as dysfunctional or exhausted. Differentiation is associated with TCR expansion, and transition from precursor to late differentiated states correlates with intratumor T cell cycling. These results provide a coherent working model for TIL origin, filiation and functional organization in primary NSCLC.

## Introduction

Lung cancer is the leading cause of cancer related death worldwide. In NSCLC (85% of lung cancers) (*1*) tumor infiltration by lymphocytes is associated with favorable survival prognosis (*2*) and a better clinical response to immune checkpoint blockade (ICB) (*3*). ICBs are thought to “re-program” CD8^+^ tumor infiltrating lymphocytes (TILs) to produce anti-tumor responses (*4*, *5*), by targeting inhibitory receptors such as PD-1, which are highly expressed by most TIL populations. How this re-programming is actually achieved, and which TIL sub-populations are targeted by ICBs are still open questions.

Most studies in the literature have distinguished early precursors and terminally differentiated CD8^+^ T cell populations (*6*, *7*) in chronic viral infections and tumors. Early precursors present characteristics of memory cells, including memory/stemness markers and regenerative capacity. They are characterized by high expression of the chemokine receptor CXCR5, which is also expressed by B cells and follicular helper T cells (TfH) (*8*, *9*). These precursors are not fully committed, and can therefore be “re-programmed” by ICB (*9*). Terminally differentiated populations are clonally expanded and express higher levels of effector markers and immune checkpoints, including PD-1, TIM-3 and CD39 (*6*, *9*). These cells share many characteristics of “exhausted” or “dysfunctional” CD8^+^ T cells. Dysfunctional TILs were initially described in chronic viral infection mouse models by poor effector function, expression of inhibitory receptors, low proliferation and distinct transcriptional and epigenetic states compared to effector or memory T cells (*10*, *11*). Several groups have recently analyzed T cell transcriptional programming in cancer, revealing a strong heterogeneity among TILs (*12*–*17*), similar to chronic viral infections, with distinct subpopulations of exhausted progenitors and terminally differentiated T cells. The idea that terminally differentiated T cells in cancer are dysfunctional, however, is still debated, especially in human tumors.

Here, we used a combination of single cell RNA and TCR sequencing (scRNA-seq and scTCR-seq) in tumors, normal tissues adjacent to the tumor (juxta-tumor), and circulating blood, to delineate the processes of CD8^+^ T cell differentiation in patients with untreated, primary NSCLC. We show that precursor, memory-like CD8^+^ TILs include 2 main populations: one is also present in the blood (circulating precursors); the other is also present in juxta-tumor tissue and bears markers of memory resident T cells (resident precursors). Both precursor subtypes differentiate into a main population of terminal effectors through a similar “transitional” stage. Terminal effectors are not observed in blood or juxta-tumor tissue, are more clonally expanded, and express signatures of exhaustion. A significant proportion of transitional and terminal effectors also express cell cycle signatures and Ki67, suggesting that clonal expansion occuring *in situ* is part of the terminal differentiation process.

## Results

### Transcriptional states of CD3^+^ tumor-infiltrating lymphocytes in NSCLC

We performed scRNA-seq on CD3^+^ T cells from 11 primary, untreated, early-stage resected NSCLC patients (Fig. 1A). For six of these patients, matched scTCR-seq information was also collected. Combined analysis of all patient samples identified a large source of variation associated with 3’ versus 5’-oriented single cell chemistries (Fig. S1A-C) that was resolved using computational methods (*18*). Rigorous assessment of intra-sample heterogeneity, including TCR, showed that integration did not over correct the sample expression profiles. After removing contaminating cell types, we collected data on 28,936 cells over the 11 patients (Fig. 1B). Uniform Manifold Approximation and Projection (UMAP) and unsupervised graph-based clustering partitioned cells into 21 clusters based on their transcriptome (Fig. 1B and C). Differential gene expression analyses (Fig.S1D, Table S1) and analyses of canonical marker genes (Fig 1D) revealed the identities of the major cell clusters. We used gene signatures (Table S2) (*17*, *19*–*24*) to reduce the impact of sparsity in the scRNA-seq values, and assign identities to different clusters, including naive, memory, regulatory, helper and effectors (Fig 1E). Among these clusters, CD4^+^ Tregs (CD4-IL32-Tregs) and CD4^+^ memory (CD4-CD69-activated memory) are the most abundant, while the most abundant CD8^+^ subsets are CD8^+^ circulating precursors (CD8-KLF2) and transitional CD8^+^ (CD8-GZMH) (Fig. 1C, Fig.S1E). We identify 10 clusters of CD4^+^ cells (including CCR8-Tregs, IL32-Tregs, GZMA-effectors, SELL-naïve, CD69-activated memory, IL7R-memory, MAL-Tregs, TNFRSF18-Tfh, SESN1-Tfh, HSPH1-memory), 7 clusters of CD8^+^ cells (including FCGR3A-effectors, KLF2-circulating precursors, GZMK-circulating precursors, XCL1-resident precursors, LAYN-terminally differentiated, GZMH-transitional, SLC4A10-MAIT), a cluster of γδ T cells (TRDC), and 3 clusters of both CD4^+^ and CD8^+^ cells: one characterized by high expression of interferon-related genes (CD4/8-ISG15) and the other two characterized by strong expression of cycling genes (CD4/8-MCM5, CD4/8-TOP2A). Analysis of immune checkpoint (ICP) molecules expression showed that CD4-CCR8 cells highly express stimulatory molecules such as TNFRSF18, while on the contrary CD8-LAYN cells show higher expression of inhibitory molecules such as HAVCR2 and LAG3 (Fig. S1F). All clusters are present in all patients, although at varying frequencies (Fig.S1G). As CD8^+^ T cells are critical players during tumor rejection, we focus our analysis on this compartment.

**Figure 1.**
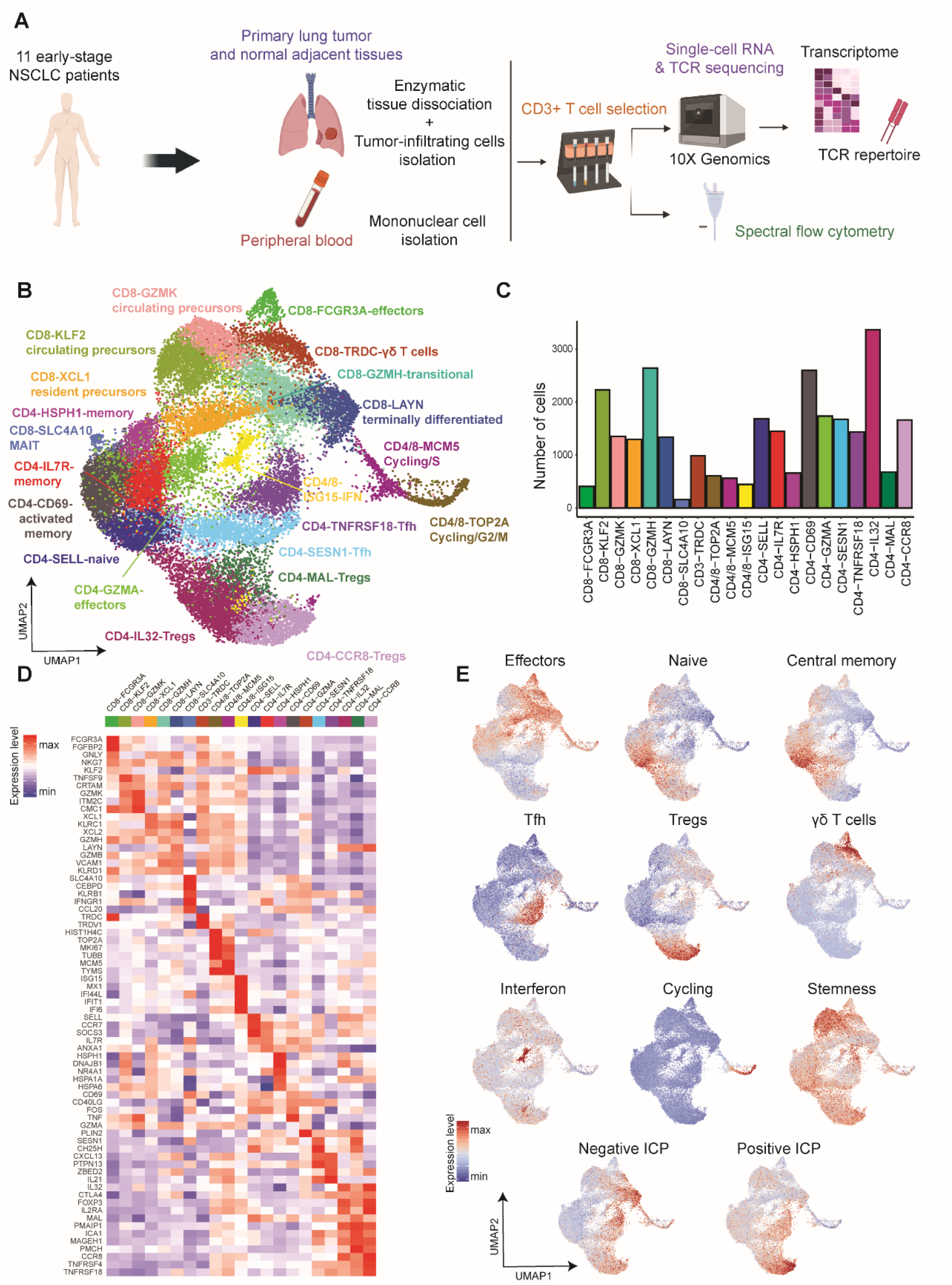
Characterization of CD3+ tumor-infiltrating lymphocytes in NSCLC. **(A)** Graphical overview of the study design. Single cell RNA and TCR sequencing was applied to tumor tissues, normal adjacent lung tissues and blood samples derived from 11 primary, untreated NSCLC patients. Spectral flow cytometry analysis was applied in an extra dataset of patients with the same clinical characteristics. **(B)** Uniform manifold approximation and projection (UMAP) of 28,936 single CD3+tumor-infiltrating T cells from 11 NSCLC patients, showing the formation of 21 main clusters, including 7 for CD8+ cells, 11 for CD4+ cells, 1 for T cells highly expressing IFN-related genes and 2 for cycling T cells. **(C)** Summary of the distribution of the number of cells contributing to each cluster. **(D)** Heatmap of expression values for the top genes with enriched expression in CD3+T cells, discovered by k-nearest neighbors’ sub-clustering. Expression values are zero-centered and scaled for each gene. **(E)** Projection of a selected set of marker genes and gene signatures, identifying T cell state. Each cell is colored based on the normalized expression.

### Memory-like precursors and terminally differentiated CD8^+^ TILs

CD8^+^ T cells segregate mainly along a horizontal axis in the UMAP, with cells expressing a stemness signature (Fig. 1E), central memory-related genes (including *IL7R, CXCR5, TCF7*, Fig. 2A) and oxidative phosphorylation signatures (also related to T cell memory, Fig. 2A) at the left end of the axis. At the right end of the axis, cells express higher levels of effector-related genes (Fig. 1E), negative immune checkpoints (Fig. 1E), including *HAVCR2* and *ENTPD1* (Fig. 2A), *TOX* (a transcription factor involved in exhaustion, Fig.2A) and a glycolysis signature (also associated with effector function, Fig. 2A). Consistent with these results, exhausted precursor (GMZK/ZNF683) and terminally exhausted (LAYN) signatures from Guo et al. (*13*) are also highly expressed on the left and right ends of this axis, respectively. Comparison of our dataset with a study on chronic viral infection in mice (*25*), shows progenitor and terminal exhausted signatures with enhanced expression on the left and right ends of this axis, respectively (Fig.S2A and B). These results suggest that CD8^+^ T cell clusters on the left of the UMAP are “memory-like precursors” (clusters CD8-GZMK, CD8-KLF2 and CD8-XCL1) and those on the right part of the UMAP (CD8-GZMH and CD8-LAYN) are more differentiated, and most likely related to late exhausted/dysfunctional cells.

**Figure 2.**
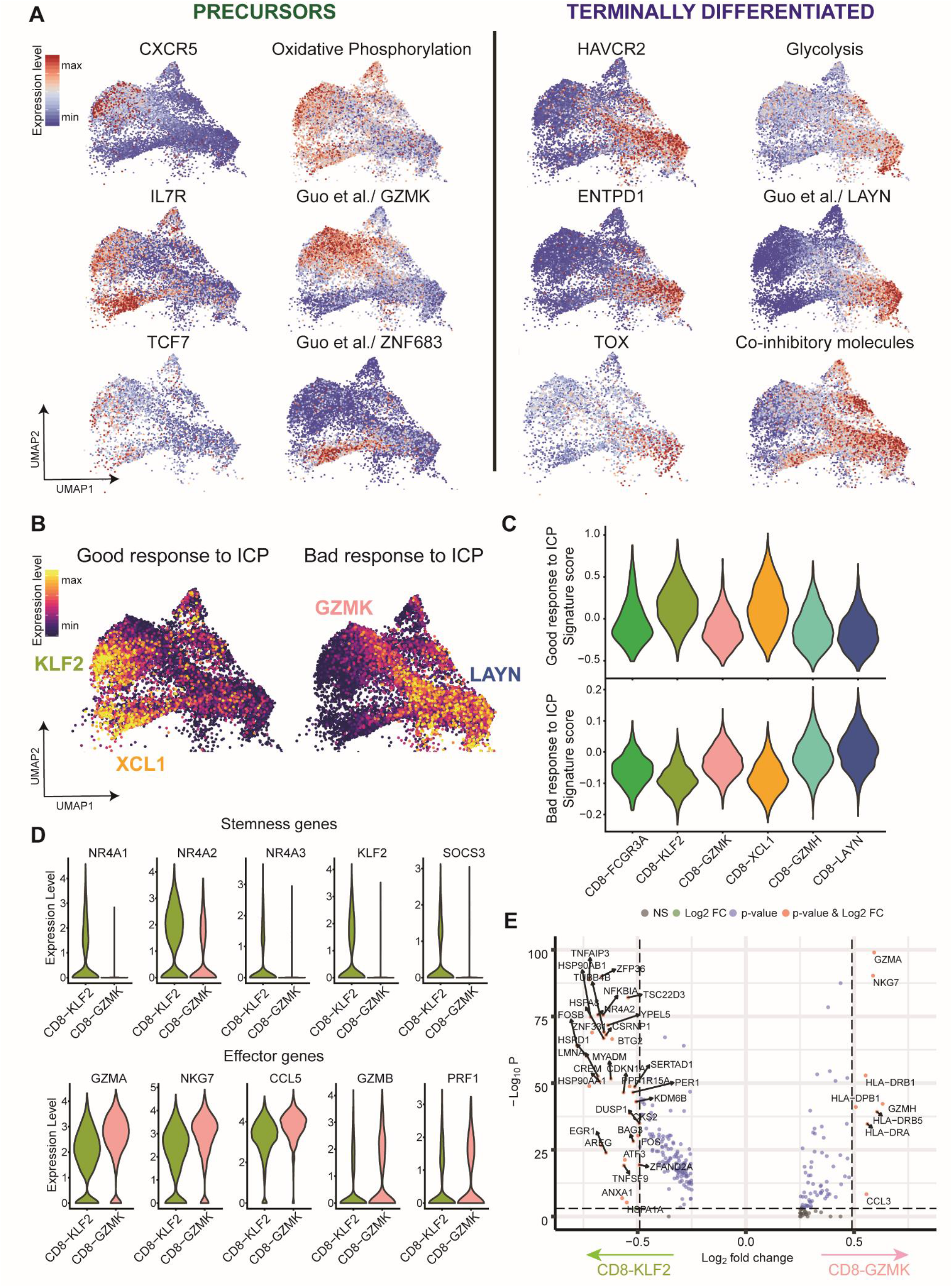
Stem-like precursors and late dysfunctional CD8+ cells. **(A)** Expression of single marker genes and gene signatures for memory-like, precursors CD8+ cells (left panel) and terminally differentiated CD8+ cells (right panel) respectively. **(B)** UMAP projection of tumor-infiltrating CD8+ cells, with each cell colored based on the gene signature score of a published study in melanoma patients. Left panel shows the expression of genes associated with therapeutic response and improved outcome to anti-PD-1 treatment (good response); right panel shows the expression of genes associated with bad response to anti-PD-1 treatment. **(C)** Violin plots of the distribution of the expression scores of the 2 gene signatures among the different CD8+ clusters. **(D)** Volcano plot of differentially expressed genes between CD8-KLF2 and CD8-GZMK clusters. Each red dot denotes an individual gene passing our p value and fold difference thresholds.

Memory-like precursors include 2 main sub-clusters, CD8-GZMK/CD8-KLF2 and CD8-XCL1, expressing high levels of the Guo et al. (*13*) GZMK and ZNF683 signatures respectively (Fig. 2A). Memory-like precursors are enriched in signatures of negative regulation of cell proliferation, while terminally differentiated cells are enriched for lymphocyte activation-related genes and adhesion integrins (Fig. S2B). We conclude that early stage NSCLC TILs include two main populations of memory-like precursors (CD8-GZMK/CD8-KFL2 and CD8-XCL1) and two populations of late, differentiated cells (CD8-GZMH and CD8-LAYN) that could correspond to terminally exhausted TILs.

### Two distinct circulating precursor subsets

Previous studies have associated precursor CD8^+^ TILs to favorable prognosis and to better response to ICB-based immunotherapies (*25*). To investigate the possible relevance of the different clusters defined here to clinical outcomes, we compared our dataset to a previous single-cell study from Sade-Feldman et al. (*17*) and applied signatures from melanoma patients responding (good response) versus not responding (bad response) to ICB. CD8-KLF2-circulating and CD8-XCL1-resident precursor clusters express higher levels of the good response signature (Fig. 2B and C). In contrast, CD8-LAYN and CD8-GZMH late clusters, and a part of CD8-GZMK early clusters, express higher levels of the poor response signature (Fig. 2B and C).

Differential analysis of genes expressed in the CD8-KLF2 and CD8-GZMK clusters reveals that cells in the latter overexpress genes related to T cell activation (HLA-DR/DP/DQ), effector related genes (KLRG1, GZMB, H, A, in addition to GZMK) and chemokines (CXCR6, CCL5) (Fig. 2D and E). The CD8-KLF2 cluster overexpresses genes related to the control of CD8^+^ T cell differentiation, including NR4A1/2/3 and KLF2 (Fig. 2D and E). These two closely related populations of CD8^+^ TILs are also present in other public datasets (Sade Feldman et al. 2018, Guo et al. 2018, Azizi et al. 2018) (*13*, *16*, *17*), including some generated on different technological platforms (SMART-Seq2/Mars-Seq) and in other tumor types, such as melanoma and breast carcinoma (Fig. S2C).

### Tissue-resident and transitional CD8^+^ populations

To better understand the different early and late TIL populations, and because one of the main driving genes among the 3 early clusters is ZNF683 (also known as Hobit, a transcription factor involved in the programming of tissue resident memory CD8^+^ T cells (*26*)), we hypothesized that some clusters could correspond to tissue resident and others to circulating cells. As shown in Fig. 3A and Fig.S3A, single gene expression of ZNF683, as well as a core tissue residency gene signature, consisting of 4 main tissue resident markers: ITGAE, ITGA1, CXCR6 and ZNF683, are all expressed at higher levels in late differentiated CD8-GZMH/CD8-LAYN and in early CD8-XCL1 clusters, as compared to the CD8-GZMK/CD8-KLF2 clusters, which are characterized by enriched expression of KLRG1 (Fig. S3A). These mRNA expression differences were validated at the protein level. Flow cytometry analysis in a set of 6 additional NSCLC patients showed that CD103^+^ CD8^+^ TILs express higher levels of markers related to T cell dysfunction, including PD-1, TIM3 and CD39, as compared to CD103^-^ TILs (Fig. 3B-D, Fig. S3B). A large fraction of CD103^+^ CD8^+^ TILs co-express T cell dysfunction markers and GZMB, while showing lower expression of KLRG1, as compared to CD103^-^ CD8^+^ TILs (Fig. 3C). These results are consistent with recent studies (*27*–*29*) that found subsets of tissue resident (CD103^+^) CD8^+^ TILs enriched in checkpoint receptors and display features of enhanced cytotoxicity and tumor reactivity, indicative of a therapeutic potential of this population.

**Figure 3.**
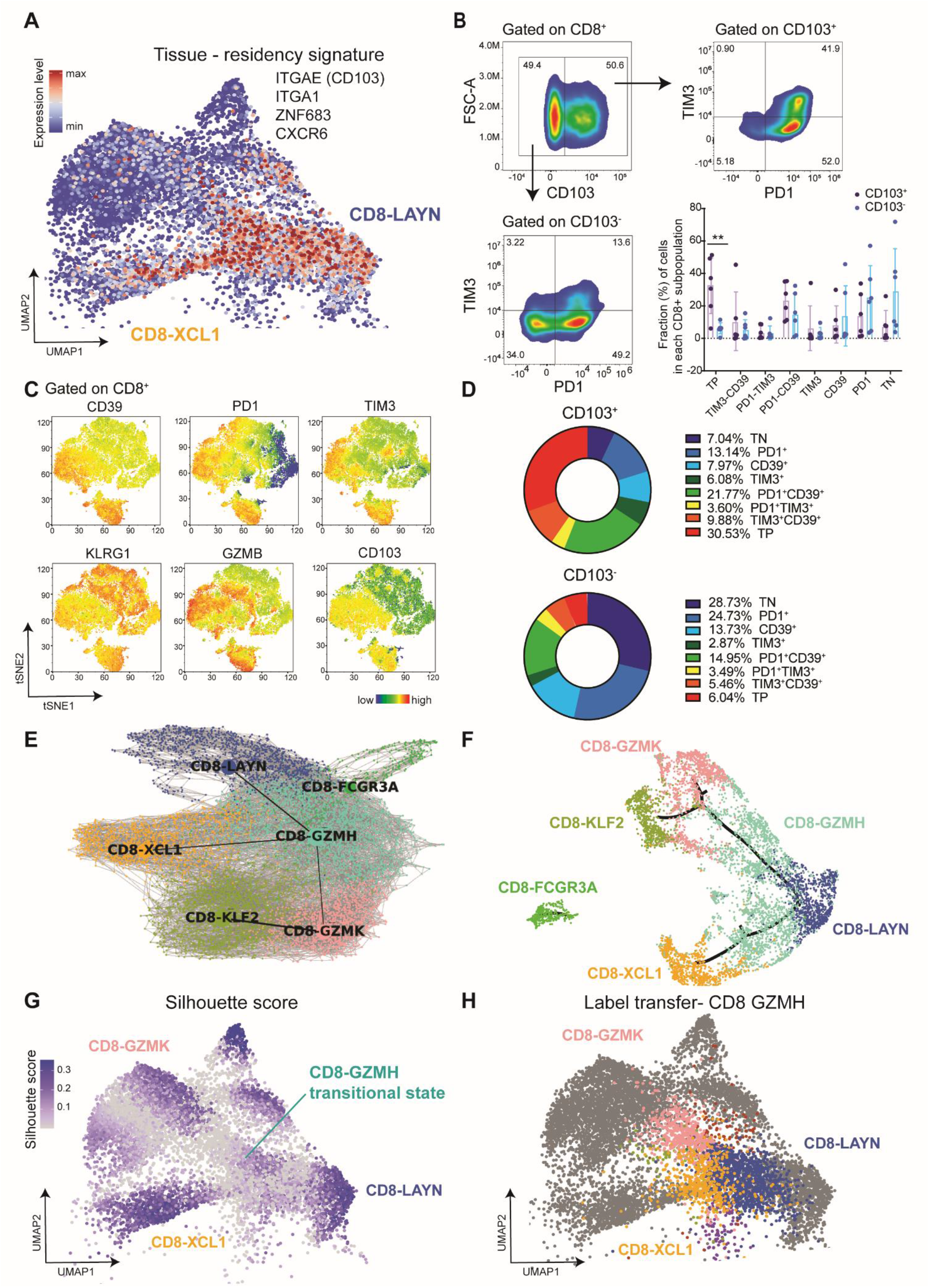
Tissue-resident and transitional CD8^+^ subsets. **(A)** UMAP projection of tumor infiltrating CD8+ cells, with each cell colored based on the relative normalized expression of a gene signature for tissue-residency, consisting of the following genes: ITGAE, ITGA1, XCL1 and ZNF683. **(B)** Flow cytometry plots of PD1 and TIM3 expression in subsets of CD103+ CD8+ and CD103-CD8+ cells. Frequency of CD103+ and CD103-cells in 8 different subsets of CD8+ tumor-infiltrating lymphocytes, defined by the expression of 3 inhibitory ICP molecules: PD-1, TIM3 and CD39. Data are representative of 6 independent experiments / NSCLC patients (**p < 0.01, two-way ANOVA test) **(C)** Representative tSNE analysis of tumor infiltrating CD8+ cells with each cell colored based on the median fluorescence intensity of each of the markers: CD103, PD1, TIM3, CD39, KLRG1 and GZMB. **(D)** Summary of the frequency of 8 different CD8+ cell subsets inside CD103+ and CD103 - CD8+ cells. **(E)** Partition-based graph abstraction of CD8+ tumor-infiltrating cells, from 11 NSCLC patients, connects circulating (GZMK, KLF2) and resident (XCL1) CD8+ precursors with terminal differentiated CD8+ (LAYN), through GZMH cluster; without any direct connection between effector CD8+ cluster (FCGR3A). **(F)** Trajectory of CD8+tumor-infiltrating cells transition state, from 11 NSCLC patients in a 2-dimensional state-space determined by Monocle v3. Each dot represents a single cell. Each color represents a different CD8+ cluster. **(G)** Visualization of the silhouette coefficient-score on the UMAP of the integrated CD8+ cells reference from 11 NSCLC tumor samples. Silhouette coefficient is calculated based on the mean intra-cluster distance and the mean of the nearest cluster distance for each cell of each cluster. Each cell is colored based on the score: 1 is the highest value, highlighted in purple and indicating robust clusters; −1 is the lower value, highlighted in grey and indicating less robust clusters; 0 is indicative of overlapping clusters. **(H)** Label transfer of cell type labels from GZMH cluster onto the integrated CD8+ cells reference from 11 NSCLC tumor samples. Colored cells are cells from CD8-GZMH cluster and grey cells are coming from the other clusters.

Analysis of cluster membership as a function of clustering resolution showed that the split between CD8-XCL1 and CD8-KFL2/GZMK occurs early, indicating the importance and robustness of the differences (Fig. S3C). Based on these results, we hypothesized that the two early, memory-like populations of CD8^+^ cells could correspond to tissue-resident (CD8-XCL1) and recent emigrants (most likely from blood, CD8-KFL2/CD8-GZMK, see below). We further speculated that the two types of precursors could both show converging differentiation into late differentiated and dysfunctional/exhausted (CD8-GZMH/CD8-LAYN) subtypes.

To test this working model, we used two different unsupervised approaches to infer continuous transitions between clusters. Statistical analysis of connectivity in the k-nearest neighbor graph of cell-cell expression similarities (PAGA) (*30*) shows that the CD8-GZMH cluster represents a transitional state between early and late clusters (Fig. 3E). Because discretizing elements in a continuum of differentiation might not be optimal, we also used pseudo-time alignment to resolve relationships between continuous populations. Monocle3 analysis revealed a converging differentiation process, consistent with the original UMAP representation and with PAGA analysis (Fig. 3F). As anticipated earlier, pseudo-time reconstitution distinguishes separate resident and circulating precursor differentiation branches (Fig. 3F). Examples of differentially expressed genes between branches are shown in Fig. S4D. With both methods, the CD8-GZMH cluster localized at the intersection between three branches: CD8-XCL1, CD8-KFL2/CD8-GZMK and CD8-LAYN, suggesting that the CD8-GZMH cluster corresponds to a transitional population of cells undergoing differentiation from early, memory-like (CD8-XCL1, CD8-KFL2/CD8-GZMK) to late, terminally differentiated (CD8-LAYN) clusters.

If the CD8-GZMH cluster indeed represents a transitional state, it should present lower clustering robustness, since cells from different clusters would enter this “state” and then differentiate into other states. We quantified robustness of clustering using the silhouette score (Fig.3G, Fig. S3E), which validates consistency by evaluating how close each cell inside a cluster is to its neighboring cells within the same cluster (high score), compared to its neighboring cells in other clusters (low score). As shown in Fig.3G, the silhouette score is lower for cells in the CD8-GZMH cluster, as compared to cells in all other CD8^+^clusters. Consistent with this analysis, label transfer, a method introduced for the integration of multiple single-cell datasets (*18*), of cells from the CD8-GZMH cluster shows even projection to the corresponding neighboring clusters, including CD8-GZMK (upper left part of the UMAP), CD8-XCL1 (lower left) and CD8-LAYN (to the right) (Fig 3H, Fig. S3F). These results, as hypothesized earlier, are consistent with early precursor CD8^+^ populations (CD8-GZMK, CD8-KLF2 and CD8-XCL1) differentiating into terminally dysfunctional CD8-LAYN cells, via a transitional CD8-GZMH state.

### Clonal sharing between clusters informs differentiation

To better understand the filiation between the different TIL clusters, we next sought to analyze their TCR repertoires. scTCR-seq for paired alpha and beta chains from 6 patients was obtained, with matched 5’ RNA profiling, for approximately 80% of the cells, including cells harboring unique or shared TCRs, indicative of clonal expansion. TCR clonotypes were called using the 10X pipeline when the analysis was restricted to one sample, and a custom algorithm when the samples were distinct but autologous (*31*, *32*). The clone size is defined as the total number of cellular barcodes associated with the same clonotype. Top clones cluster identities for each patient are shown in Fig. S4A.

To investigate the extent of clonal expansion in different clusters, we first represented individual cells colored according to the size of their TCR clones. As shown in Fig. 4A and B, larger clones are found within CD8^+^ T cell clusters as opposed to CD4^+^, and more specifically in late differentiated (CD8-GZMH and CD8-LAYN) clusters. This pattern was conserved across patients (Fig. 4B, lower panel), as measured using either a published expansion index (*33*) or simply by the mean number of cells per TCR clonotype. When visualizing all cells from individual TCR clones in the UMAP we found that in most cases TCR clones are confined to specific clusters (examples in Fig. 4C).

**Figure 4.**
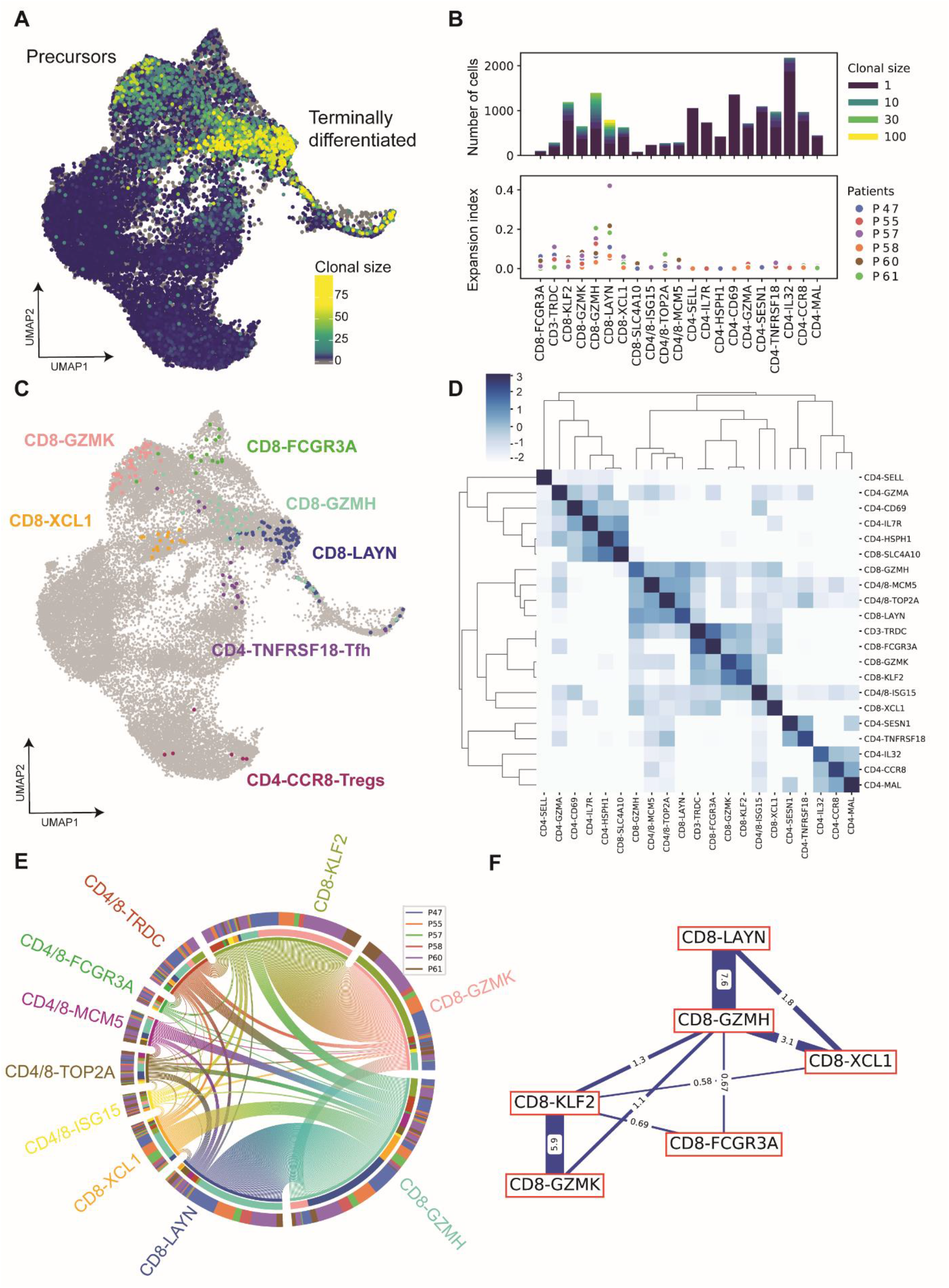
Clonal expansion and TCR sharing in clusters. **(A)** Visualization of the clonal size on a UMAP embedding on the integrated 11 tumors reference. **(B)** Upper panel: Quantification of clonal size per cluster identities; Lower panel: Expansion index computed for each cluster and each patient. (C) Selected examples of cluster-specific clones from P60. **(D)** Normalized heatmap of clonal sharing probabilities (as defined in the subsection TCR analysis of the methods). **(E)** Circos plot of clonal sharing between clusters. Each line represents one shared clone with the most frequent sharing. **(F)** Scheme of sharing similarities between CD8 clusters as computed in Fig.4D.

To quantify if there is a link between individual TCRs and specific clusters, we calculated the conditional probability for a particular TCR observed in one cluster to be also observed in the same or other clusters. Unsupervised clustering of these probabilities is shown in Fig. 4D. The probability that the same TCR is found in cells from the same clusters is consistently higher than the probability to find the same TCR in cells from different clusters (dark blue diagonal). Nevertheless, there is an observable degree of TCR sharing across clusters and the probability to find the same TCRs in two different clusters is not zero. As illustrated by the light blue cases in Fig. 4D, “sharing” of TCRs between clusters occurs, and is preferential between clusters with similar functional abilities. For example, the sub-clusters of CD4-Tregs share more TCRs among them than they share with CD4-Tfh or CD4-memory T cell clusters. Among the CD8^+^ T cell clusters, increased sharing occurs between circulating precursor memory-like clusters (CD8-GZMK, CD8-KLF2) and between differentiated clusters (CD8-LAYN, CD8-GZMH). The CD4/8-ISG15 (enriched for interferon-related genes) and the cycling clusters show increased sharing with early memory-like and late differentiated clusters, respectively. We conclude that while high intra-cluster TCR sharing highlights the relevance of our clustering strategy, inter-cluster TCR sharing can be used to infer transitions of cells between clusters: higher TCR sharing between two clusters would indicate dynamic transitions of cells between the two transcriptional states. We conclude that inter-cluster TCR sharing can be used to infer a biologically relevant measure of proximity between clusters that can indicate either a dynamic transition between two transcriptional states or a static similarity not fully captured by transcriptome-based clustering methods.

To investigate potential transitions between clusters based on TCR sharing in more detail, we next represented all the TCRs shared between two clusters in a “circos” representation (Fig.4E), where each line represents a single TCR clonotype. This representation illustrates that the two main non-resident precursor memory-like clusters (CD8-GZMK and CD8-KLF2) and the two main late differentiated (CD8-GZMH and CD8-LAYN) clusters share numerous TCRs. We also computed the transition index using the STARTRAC method (*33*), as the likeliness of a cluster to share clones (Fig. S4B) and we also see that CD8-LAYN and CD8-GZMH are the clusters which are most likely to have TCR sharing. The circos plot also shows that GZMH displays higher clonal sharing with all the rest of the clusters (Fig. 4E, Fig. S4C). This result was reproducible between patients. The CD8-GZMH cluster shares over half of its shared TCRs with the CD8-LAYN cluster, while the other half is shared with other CD8^+^ T cell clusters (Fig. S4C). Supervised representation of the computed probabilities of sharing TCRs among clusters (Fig. 4F), shows that the CD8-GZMH cluster could represent a central hub, sharing TCRs with both early, memory-like and late, differentiated clusters. The high levels of TCR sharing between clusters, was confirmed by analysis of the number of T cells from each cluster for shared clones. As shown in Fig. S4D, “equilibrated” sharing between clusters (when the same TCR is present in 3 or more cells from each cluster) is only seeing between CD8-GZMH/CD8-LAYN and CD8-GZMK/CD8-KFL2 (and to a lower extent between XCL1/GZMH). Equilibrated sharing suggests active transitions between clusters. As proposed below, TCR sharing analysis is consistent with the GZMH cluster representing a transitional state between the two main populations of memory-like precursors (CD8-KLF2/CD8-GZMK and CD8-XCL1) and the late differentiated cells present in the CD8-LAYN cluster.

### Terminally differentiated, not memory-like, CD8^+^ T cells are in cell cycle

The question of the cycling activity of early memory-like vs. late terminally differentiated TILs is still a matter of debate (*34*). As T cells require TCR signaling to enter the cell cycle, clones identified to be actively cycling intratumorally strongly suggest a local (intratumor) source of antigen. Because cell cycle-related genes are expressed at high levels, cycling cells cluster independently in scRNA-seq analyses. These clusters, however, may include cells that also bear underlying signatures from other clusters, which are in some way “masked” by highly expressed cell cycle genes. We took two independent approaches to investigate if the cells in the cycling cluster are related to other particular clusters.

First, we focused on the transcriptomic dataset. Clustering of the infiltrating T cells in 11 NSCLC patients shows two different cycling populations, one in G2/M phase (CD4/8-TOP2A) and the other one in S phase (CD4/8-MCM5) (Fig.S5A). To better characterize these clusters, we used label transfer to interrogate the “second best” cluster to each cycling cell. Because the IFN-related cluster (CD4/8 ISG15) also includes CD4^+^ and CD8^+^ T cells, and because interferon-stimulated genes (ISGs) are also highly expressed and can drive independent clustering of cells related to other clusters, we included this cluster in the same label transfer analysis. Label transfer of CD8^+^ cells from the CD4/8 ISG15 cluster, re-cluster mainly to CD8-GZMK and CD8-GZMH clusters (Fig. 5A and B). Label transfer of CD8^+^ T cells from the cycling clusters, results in almost exclusive re-clustering to late differentiated effector clusters CD8-GZMH and CD8-LAYN (Fig. 5A and B). Virtually no cells from the cycling clusters are re-attributed to progenitor CD8^+^ T cell populations, suggesting that cycling cells are transcriptionally closer to late differentiated T cells.

**Figure 5.**
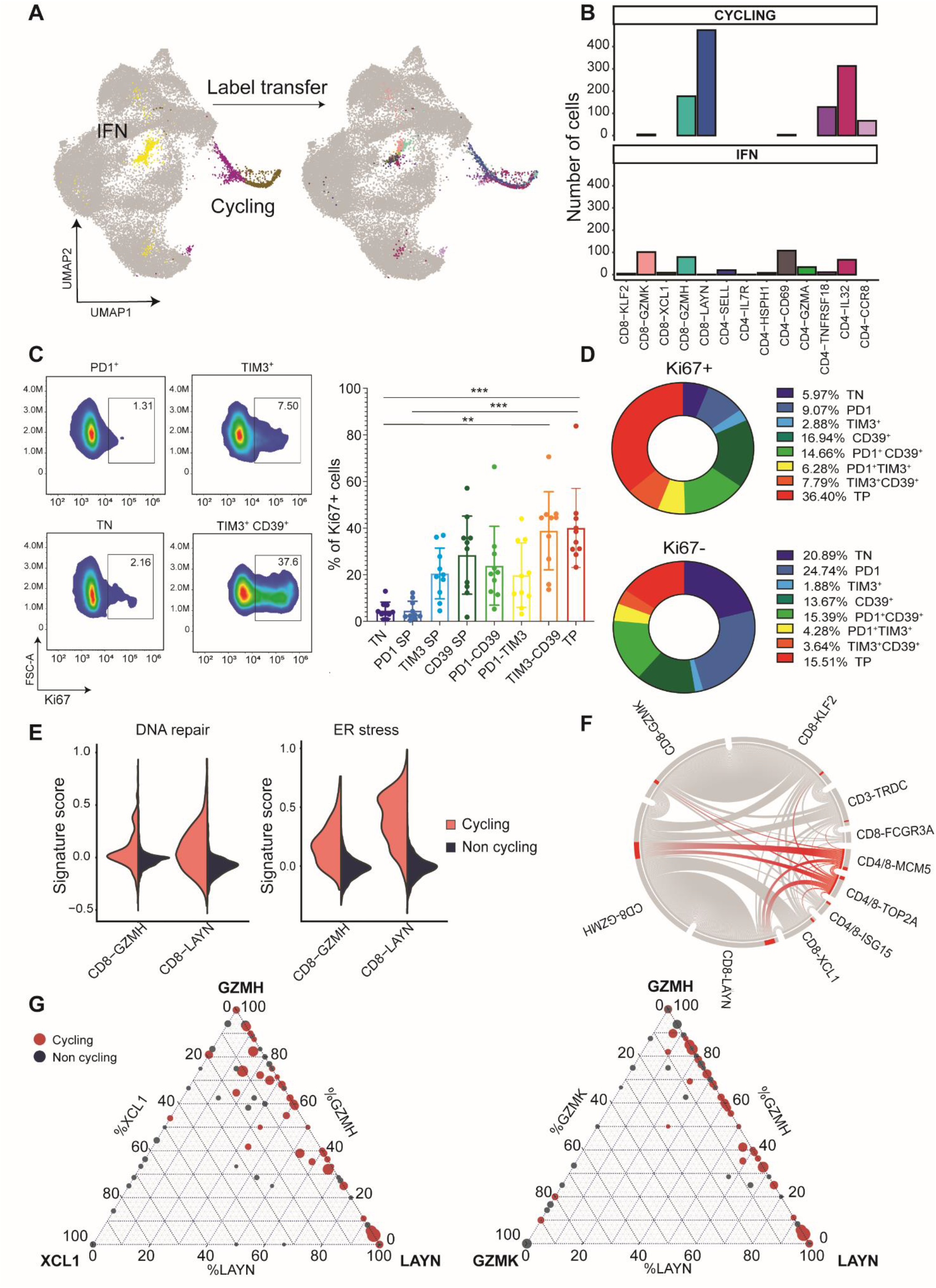
Late dysfunctional CD8^+^ cells are high proliferative. **(A)** Label transfer from the two cycling clusters (CD4/8-TOP2A and CD4/8-MCM5) and the IFN-cluster (CD4/8-ISG15) onto the integrated CD8^+^ cells reference from 11 NSCLC tumor samples. **(B)** Distribution of the number of cells label transferred from the cycling cluster (top part) and the IFN^high^ cluster (bottom part) into the CD3^+^ T cell clusters. **(C)** Representative flow cytometry plots of Ki67^+^ population in subsets of triple-negative (TN), PD1+, TIM3+ and TIM3+ CD39+ of CD8+ cells. Quantification of the frequency of Ki67+ cell in subsets of triple positive (TP), CD39+ TIM3+, PD-1+ CD39+, PD-1+ TIM3+, PD-1+, TIM3+, CD39+ and triple negative (TN) CD8+ cells, based on the expression of the 3 exhaustion markers: PD-1, TIM3 and CD39. Data are representative of 10 independent experiments / NSCLC patients (mean ± SEM) (***p<0.001, **p < 0.01; two-way ANOVA test) **(D)** Summary of the frequency of 8 different CD8^+^ cell subsets (based on the expression of PD-1, TIM3, CD39; triple positive (TP), CD39^+^ TIM3^+^, PD-1^+^ CD39^+^, PD-1^+^ TIM3^+^, PD-1^+^, TIM3^+^, CD39^+^, triple negative (TN) inside the populations of Ki67^+^ and CD103^-^ CD8^+^ cells. **(E)** Violin plots show the distribution of expression of genes associated to DNA repair and ER stress between cycling and non-cycling subsets of CD8-LAYN and CD8-GZMH clusters. **(F)** Circos plot of clonal sharing between the two cycling clusters (CD4/8-TOP2A and CD4/8-MCM5) and the rest of CD8^+^ clusters (highlighted in red); each line represents one shared clone. **(G)** Quantification of each cluster contribution to shared clones. Each dot corresponds to a shared clone between the 3 clusters: CD8-GZMH, CD8-LAYN, CD8-GZMK (left panel) or CD8-GZMH, CD8-LAYN, CD8-XCL1 (right panel). Dots highlighted in red correspond to clones that are shared with the cycling clusters.

We validated these results using flow cytometry analysis in freshly isolated TILs from 10 additional NSCLC patients. As shown in Fig. 5C and D, CD8^+^ cells co-expressing PD1, TIM3 and CD39 (triple-positive, TP) are also Ki67^+^. CD8^+^ T cells negative for the 3 inhibitory ICP markers or expressing only PD1, show very low Ki67 labeling (Fig. 5C and D). tSNE visualization of cytometry data shows that a tissue-resident (CD103^+^) subset of these proliferative, triple-positive CD8^+^ cells also expresses PD-L1, suggesting that this subpopulation could play an important role in ICP response (Fig.S5B).

The CD8-LAYN subset has previously been associated with exhausted/dysfunctional CD8^+^ T cells (*13*, *33*, *35*, *36*). Among cycling cells, the ones transcriptionally related to the CD8-LAYN cluster by label transfer also express higher levels of DNA damage repair genes (Fig. S5C). These cells also express higher levels of gene signatures for DNA damage repair and stress (Fig. 5E), as well as of the terminal differentiated LAYN signature (*13*) (Fig. S5D), as compared to cycling cells re-attributed to the CD8-GZMH cluster. Analysis of the Guo et al. dataset (*13*) for the cycling signature shows that the LAYN and the cycling populations also overlap (Fig. S5E), suggesting that late differentiating cells, and not early progenitors, are the main cycling T cells in NSCLC TILs.

Next, we reasoned that if late differentiating cells cycle, clonally amplified TCRs in these clusters should also be found in cycling T cells. Circos representation shows that the TCRs expressed in cycling cells are found preferentially in cells from late differentiated effector CD8-GZMH and CD8-LAYN clusters (Fig. 5F). As shown in Fig. S5F, preferential TCR sharing of cycling cells with late differentiated clusters is not uniquely due to the larger size of these late clones. Indeed, CD8-GZMK/CD8-KLF2 clones of over 20 cells still share very few TCRs with the cycling cluster, as compared to CD8-GZMH/CD8-LAYN clones of similar sizes. Increased sharing can also be visualized in the sharing probability graph shown in Fig. 4D. Therefore, cycling cells share TCRs preferentially with late effectors, as compared to early memory-like precursors.

The results presented thus far are consistent with the possibility that transition between CD8-GZMH and LAYN clusters occurs while cells divide. To test this hypothesis, we measured the frequency of cycling cells among clones shared between the different clusters. As shown in Fig. 5G, the majority of clones that include GZMH and LAYN cells, also have TCRs in the cycling clusters (central dots in the right panel), as compared to fewer central cells in the two other panels. Fig. S5G, uses a similar representation to show that the clones that include TCRs from the GZMH, LAYN, and cycling clusters, frequently also include TCRs from the CD8-XCL1 cluster (as compared to also including TCRs from the GZMK cluster).

These results suggest that the preferential pathway toward terminal differentiation in tumors originates in the CD8-XCL1 cluster, and transitions through CD8-GZMH to CD8-LAYN while cell divide and clonally expand. This model is consistent with these cells being chronically stimulated, and with recent results in mice (*37*) chronic viral models.

### Resident versus circulating origin and tissue distribution of TILs

To further investigate the tissue *versus* blood origins of the different TIL populations described so far, we analyzed scRNA and TCR sequencing in TILs from blood and juxta-tumor tissue. In 4 of the previously described patients, we also obtained blood samples, and in 2, juxta-tumor tissue. scRNA-seq and scTCR-seq in all 4 patients resulted in 54,247 validated cells, with ~ 80% RNA and TCR coupled data. After removing contaminating cells and integrating the 3 different tissues, we used as a reference the clusters having been identified before in the 11 different NSCLC tumor samples (as shown in Fig.1B).

By integrating cells from the 3 different tissues in these patients and using label transfer to map the populations from blood and juxta-tumor into the tumor reference, we quantified the proportions of the different populations in the different tissues (Fig.6A-C). This allowed us to focus on conserved patterns between our tumor reference and the other tissues, and minimize over-smoothing induced by integrating data from different tissues (Fig. S6A). The CD4-SELL cluster (naive T cells), contains mostly cells from blood, whereas the CD8-LAYN cluster (terminally differentiated), contains almost exclusively cells from tumors (Fig. 6A-C, Fig. S6B). This result is consistent with flow cytometry analysis in 4 additional NSCLC patients, showing that TIM3^+^ CD39^+^ CD8^+^ cells are enriched in the tumor samples, as compared to blood samples, which contain mainly single positive PD1^+^, or TN CD8^+^ cells (Fig. S6C).

**Figure 6.**
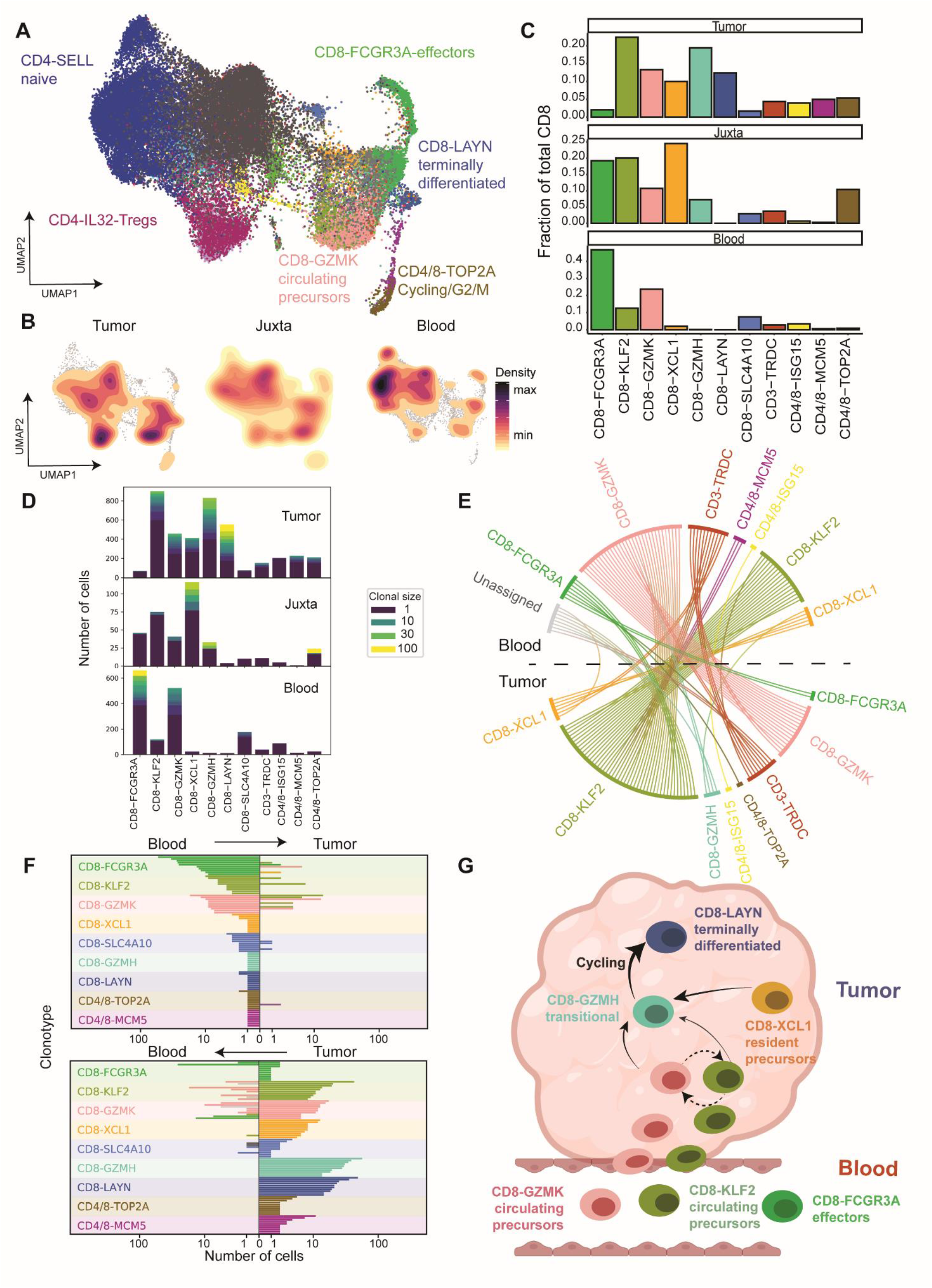
Tissue-specific features - integrative model. **(A)** UMAP representation of the integrated cells from 17 samples from tumor, juxta-tumor and blood. **(B)** UMAP representation of cell densities split by the 3 tissues. **(C)** Scaled proportions of cells per clusters across the 3 tissues. **(D)** Quantification of clonal expansion for each cluster across the 3 tissues. **(E)** Circos plots of clonal sharing between Tumor and Blood cells. Each line represents one shared clone with the most frequent sharing. **(F)** Top 10 shared clones of blood (upper panel) and tumor (bottom panel) being shared with tumor and blood respectively for each CD8 cluster. **(G)** Summary scheme of CD8 circulation and differentiation in NSCLC. Graph created using BioRender software (BioRender.com)

The CD8-XCL1 cluster (resident precursors) contains mostly cells from tumor and juxta-tumor tissue (not from blood), indicating that it is indeed a tissue-specific population, while cells in the CD8-GZMK and CD8-KFL2 clusters are also present in blood. These results are consistent with the proposed dual origin of memory-like progenitors: CD8-KLF2/CD8-GZMK cells originate from blood, and would represent circulating precursors, while CD8-XCL1 cells absent from blood, but in juxta-tumor tissue, would represent tissue-resident precursors (Fig. 6B and C). The most abundant CD8^+^ T cell cluster in blood is CD8-FCGR3A, which are also present in juxta tumor samples, but are very rare in tumors (Fig. 6B and C). Also consistent with our model, CD8-GZMH (transitional) and CD8-LAYN (terminal) clusters are phenotypes acquired in tissue (Fig. 6B). The absence of CD8-LAYN cells in blood and juxta-tumor is also consistent with CD8-GZMH cells becoming CD8-LAYN only in the tumor microenvironment.

Finally, we used scTCR information to further test this model. Most TCR expanded clones in the 3 tissues are present within CD8^+^ T cell populations (Fig 6D). The clusters with the most expansion, however, are distinct in each tissue: CD8-LAYN in tumors, CD8-XCL1 in juxta-tumor and CD8-FCGR3A (together with the CD8-GZMK/CD8-KLF2) in blood (Fig. 6D). Analysis of shared TCR clonotypes present in tumor and blood (Fig. 6E), shows extensive TCR sharing between CD8-GZMK and CD8-KLF2 clusters in the two locations. Many expanded CD8-GZMK TCRs in blood are expressed in CD8-KLF2 cells in tumor, suggesting that CD8-KLF2 TILs may be derived from CD8-GZMK blood cells.

To further investigate possible filiations between clusters in blood and tumor, we analyzed the top 20 TCR clones per cluster in the blood (Fig. 6F, upper panel) or in the tumor (Fig. 6F, bottom panel) for their transcriptional programing and TCR numbers in the other tissue (blood for tumor and tumor for blood). The top 20 TCR clonotypes from late differentiated tumor clusters (CD8-GZMH and CD8-LAYN) were not observed in the blood, consistent with these clones expanding intratumorally. Among the top 20 CD8-XCL1 TCR clonotypes from the tumor, only one was also found in blood, where it has CD8-GZMK transcriptional programing. This result is consistent with most CD8-XCL1 clones not coming from blood, but having tissue-resident origin. Among the top 20 tumor clonotypes in the CD8-KLF2 and CD8-GZMK clusters, all are found at significant frequencies in blood. However, while top tumor CD8-GZMK clonotypes are also mapped to CD8-GZMK cells in blood, top tumor CD8-KLF2 clonotypes are also found in CD8-GZMK clustered cells in blood (as suggested by the circos analysis, Fig. 6E). Analysis of the top 20 clonotypes per cluster from the blood yields consistent results. First, most expanded clones in blood are in the CD8-GZMK cluster, and most of these clones are also found in the tumor, where they display transcriptomic re-programing corresponding to both CD8-KLF2 and CD8-GZMH clusters, mainly. This result suggests that TCR expanded CD8-GZMK blood CD8^+^ T cells are the main blood precursors for tumor infiltration. They also suggest that after infiltration, blood TCR expanded CD8-GZMK precursors are re-programmed to other phenotypes including CD8-KLF2 and CD8-GZMH.

## Discussion

Here, we used scRNA- and scTCR-seq to analyze CD8^+^ TILs, juxta-tumor tissue and blood in early-stage resected NSCLC patients. Based on integration of transcriptomic programing with TCR in the 3 tissues, we propose an integrative working model for TIL origin, filiation and functional organization in primary NSCLC (Fig. 6G).

The most abundant and clonally expanded population of CD8^+^ T cell effectors in blood (CD8-FCGR3A) are very rare in tumors (as are the clonal TCRs they express), indicating that these cells do not infiltrate tumors effectively (if the cells entered tumors and changed phenotypes, their TCRs would still be present in other clusters). CD8-GZMK cells, in contrast, are abundant and clonally expanded in both blood and in tumors, and share similar TCRs, suggesting that they represent the main blood precursor for TILs. A significant proportion of the TCRs present in the CD8-GZMK cluster are also present in tumors in a second cluster of memory-like cells, CD8-KLF2, suggesting that these two populations are in dynamic equilibrium. Although trajectory reconstitution suggests that these two populations can differentiate into transitional CD8-GZMH cells, TCR sharing between the two is relatively low as compared to the sharing between CD8-XCL1 and CD8-GMZH. This result suggests that the transition between CD8-GZMK/CD8-KLF2 and CD8-GZMH is slow or inefficient. The other main population of early, memory-like, CD8^+^ T cells are in the CD8-XCL1 cluster. These cells express tissue resident signatures and markers, and are absent from blood, suggesting their tissue resident origin. These cells share numerous expanded TCRs with CD8-GZMH cells, suggesting that they represent the main source of GZMH transitional cells.

The effector-like, late CD8^+^ TILs are divided into 2 clusters, CD8-GZMH and CD8-LAYN. Cells in the CD8-GZMH cluster show limited robustness in clustering and share numerous clonal TCRs with all other CD8^+^ T cell clusters, indicating that early-memory like cells (circulating or tissue resident) transit through this stage before entering late terminally differentiated, most likely dysfunctional states (CD8-LAYN). Cells in this cluster belong to the larger TCR clones, most of which are shared with the CD8-GZMH cluster. Consistent with the idea that transition between these two late clusters requires cell divisions, a significant proportion (~40%) of these late stage cells show strong cell cycle signatures and are labeled by Ki67 antibodies. Intratumor expansion of these cells, together with expression of T cell activation markers suggest that the cells recognize antigens intratumorally (whether tumor-specific or not). Since tissue-resident CD8-XCL1 cluster cells share most TCRs with the CD8-GZMH/CD8-LAYN clusters, we would hypothesize that tumor antigen-specific T cell derive mostly from tissue-resident memory populations (rather than from clones recently stimulated by antigen in the tumor-draining lymph nodes). Indeed, the highly expanded TCRs present in blood (mainly in CD8-GZMK cluster cells) remain in early/memory-like clusters, and are rare in late clusters, suggesting rather inefficient intratumor progression from blood precursors to terminal differentiated cells.

Whether cells in the CD8-LAYN cluster are active effectors or dysfunctional/terminally exhausted was not investigated in this study. Several recent papers provide partial evidence in each direction, depending on the types of tumors and markers used (*14*, *19*, *28*, *36*–*38*). Here, we show that the CD8-LAYN cluster express high levels of a “bad response” signature to ICP in melanoma. We also found that cycling cells with LAYN-signature expression also highly express ER stress and DNA repair signatures, as compared to both non-cycling cells from the same cluster and to cycling cells from the CD8-GZMH cluster (Fig.5E). These results are consistent with cycling LAYN cells being exhausted/dysfunction, rather than actively involved in tumor rejection.

In contrast to CD8-LAYN cells, very few memory-like CD8^+^ T cells were found to bear cell cycle signatures. In mice, several studies (and our own unpublished results) show that progenitor, memory-like TILs cycle (*36*–*38*), while the human results are more controversial. Recent papers suggest that TILs in early-dysfunctional state in melanoma cycle (*14*), while other studies in NSCLC are consistent with our results, showing that more terminally differentiated cells, highly expressing inhibitory ICP molecules cycle more than early progenitors (*12*). The working model for TIL infiltration and differentiation proposed here would predict that TCR expansion in the CD8-LAYN cluster (most likely “exhausted” or ”dysfunctional” T cells) will not be of good prognosis for clinical response to ICP blockers. Indeed, abundant literature suggests that terminally differentiated T cells cannot be re-programmed to reject tumors. Among early, memory-like, potentially reprogrammable CD8 clusters, our results suggest that TCR expansion in CD8-XCL1, rather than CD8-KLF2/CD8-GZMK, cluster cells could be of good prognosis for response to ICB. Future scRNA-seq studies in NSCLC patients responding or not to ICB will test this hypothesis.

## Materials and Methods

### Software versions

Data were collected using Cell Ranger software (10X Genomics) v.2.2.032, and analyzed using R v.3.5.1, and the following packages and versions in R for analysis: Seurat v3.1.1, ENHANCE v1.0.0, DroptletUtils v1.8, clustree v0.4.1, cluster v2.1.0 Two-dimensional gene expression maps were generated using coordinates from the UMAP algorithm^39^ using the R package uwot v0.1.3 implementation. Figures were produced using the following packages and versions in R: RColorBrewer v1.1-2; pheatmap v1.0.12; ggplot v3.2.0; ggsignif v0.6.0.

### Human specimens

Eleven patients, who were pathologically diagnosed with NSCLC were enrolled in this study, including 10 adenocarcinoma and one squamous-cell carcinoma patients. All patients were untreated, in early disease stage. Tumor tissue samples were obtained from all 11 NSCLC resected patients. For 4 of them paired peripheral blood was also collected and analyzed. Among those 4, for 2 of the patients, matched adjacent normal lung tissue was collected and analyzed. All samples were collected from the Institute Mutualiste Montsouris, under a dedicated protocol for lung cancer specimens approved by the French Ethics and Informatics Commission (EUdract 2017-A03081-52). All patients in this study provided written informed consent for sample collection and data analysis.

Tumor tissue and adjacent normal lung tissue were obtained from surgical specimens, after macroscopic examination of the tissue by a pathologist. Tissue samples were stored in C0_2_-independent medium (Invitrogen) with 10% human serum and transferred within 1-hour post-surgery to the research institute. For each specimen, a fragment was formalin-fixed, and paraffin embedded for histology and immunohistochemistry.

### Tissue dissociation

Tumor and adjacent normal lung tissue samples were gently cut in approximately 1mm3 pieces. Tissues were digested enzymatically, by an incubation of 20-40 minutes, based on the size of the tissue, at 37°C in agitation, in C0_2_-independent medium (Invitrogen) with Collagenase I (2mg/ml, Sigma-Aldrich), Hyaluronidase (2mg/ml, Sigma-Aldrich) and DNase (25μg/ml, Sigma-Aldrich). The tissue pieces were gently triturated with a 20 ml syringe plunger on a 40 μ m cell strainer (BD) in 1X PBS (Invitrogen) with 1% fetal bovine serum (FBS) and 2mM EDTA (Gibco) until uniform cell suspensions were obtained. The suspended cells were subsequently centrifuged for 10 minutes at 400g.

### Tumor-infiltrating lymphocytes isolation

Tumor-infiltrating lymphocytes were isolated using Ficoll-Paque PLUS solution (Sigma-Aldrich). After tissue digestion, cells are resuspended in C0_2_-independent medium and layered onto Ficoll-Paque PLUS solution. Subsequently cells are centrifuged for 20 minutes at room temperature at 400 rcf without breaks. After centrifugation tumor-infiltrating lymphocytes were carefully transferred to a new tube and washed with 1X PBS with 1% FBS and 2mM EDTA (Gibco).

### Peripheral blood mononuclear cells isolation

Peripheral blood mononuclear cells (PBMCs) were isolated using Ficoll-Paque PLUS solution (Sigma-Aldrich). Fresh peripheral blood was collected before the surgery in EDTA anticoagulant tubes. Briefly 8 ml of fresh peripheral blood were resuspended in 1X PBS (Invitrogen) with 1% FBS and 2mM EDTA (Gibco) and the layered onto Ficoll-Paque PLUS solution. Subsequently cells are centrifuged for 20 minutes at room temperature at 2000 rpm without breaks. After centrifugation tumor-infiltrating lymphocytes were carefully transferred to a new tube and washed with 1X PBS with 1% FBS and 2mM EDTA (Gibco).

### CD3^+^ T cell isolation and purification

CD3^+^ T cells from all samples were enriched using magnetic positive selection; CD3 microbeads and MACS separation (Miltenyi). Subsequently dead cells and debris was removed, following manufacturer instructions (Dead cell removal kit; Debris removal kit; Miltenyi Biotec) resulting in ~80% of purity and CD3^+^ T cells were resuspended in 1X PBS with 0.04% BSA. Cell numbers and viability were measured using a Countess II Automated Cell Counter (Thermo Fisher Scientific) as well as classical hemocytometer and trypan blue.

### Single-Cell RNA sequencing and TCR (VDJ) profiling

Single-cell suspensions were loaded onto a Chromium Single Cell Chip (10X Genomics) according to the manufacturer’s instructions for co-encapsulation with barcoded Gel Beads at a target capture rate of 5000 −10.000 individual cells per sample, based on the initial number of cells per sample. Referring to the blood samples the target capture rate was 10.000 individual cells, while for tumor tissue and normal adjacent lung tissue samples it was diverse between 5.000 – 10.000 individual cells, due to the diverse cell number per patient’s tissue. Captured mRNA was barcoded during cDNA synthesis and converted into pooled single-cell RNA-seq libraries for Illumina sequencing using the Chromium Single Cell 3’ Solution (10X Genomics) according to the manufacturer’s instructions. All samples for a given donor (blood, tumor and normal adjacent lung tissue) were processed simultaneously with the Chromium Controller (10X Genomics) and the resulting libraries were prepared in parallel in a single batch. After Gel Bead-in-Emulsion reverse transcription (GEM-RT) reaction and clean-up, a total of 14 cycles of PCR amplification were performed to obtain sufficient cDNAs used for both RNA-seq library generation and TCR V(D)J targeted enrichment followed by V(D)J library generation. TCR V(D)J enrichment was done per manufacturer’s user guide using Chromium Single Cell V(D)J Enrichment Kit, Human T cell (10X Genomics). cDNA before and after TCR enrichment was profiled using both Qubit (Thermofischer scientific) and Bioanalyzer High Sensitivity DNA kit (Agilent Technologies). Libraries for RNA-seq and V(D)J were prepared following the manufacturer’s user guide (10X Genomics), then profiled using Kapa Library Quantification kit (Kapa Biosystems) and quantified with Qubit (Thermo Fisher Scientific). Single-cell RNA-seq libraries were sequenced with NovaSeq system (Illumina). Single-cell TCR V(D)J libraries were sequenced with HiSeq2500 machine (Illumina). All sequencing was done according to the manufacturer’s specification (10X Genomics).

### Flow cytometry and antibodies

Flow cytometry analysis was performed in a dataset of 10 additional untreated NSCLC patients, in early disease stage. For the isolation of lymphocytes from tumor tissue and normal adjacent lung tissues, as well as of PBMCs, the same steps pre-described above were followed. For negative selection of CD3^+^ T cells we used the Pan T cell Isolation Kit (Miltenyi), that leads to approximately 80% purity. All cells were then stained in 1X PBS buffer with Zombie NIR fixable viability kit (#B262784, Biolegend) according to the manufacturer instructions, using a combination of monoclonal antibodies from the following: CD3-AF532 (1:50, clone UCHT1, #2067979, Thermofischer Scientific), CD8-BV510 (1:50, clone 2ST8SH7, #9259590, BD Biosciences), PD1-BV421 (1:40, clone EH12.2H7, #B268454, Biolegend), KLRG1-PE-Cy7 (1:40, clone SA231A2, #B256810, Biolegend), TIM3-BV786 (1:40, clone 7D3, #9259393, BD Biosciences), CD39-APC (1:50, clone eBioA1, #2071264, Thermofischer Scientific), PD-L1-APC-R700 (1:40, clone M1H1, #9010854, BD Biosciences), VCAM1-BB515 (1:20, clone S1-10C9, #9093693, BD Biosciences), CD103-PerCP-Cy5.5 (1:50, clone Ber-ACT8, #B246047, Biolegend). Intracellular staining was performed using the eBioscience™ intracellular fixation & permeabilization buffer set (Thermofischer Scientific) according to the manufacturer’s guidelines, using a combination of the following monoclonal antibodies: GZMB-PE(1:20, clone QA16A02, #372207, BD Biosciences), Ki67-BV711(1:20, clone Ki67, #B291484, Biolegend). Samples were analyzed using a Cytek^®^Aurora spectral flow cytometer. Gating was applied by using monocolors and FMO controls. Data analysis was performed using FlowJo v.10.6.1.

### Statistical analysis

All flow cytometry experiments were analyzed using Prism 8 (GraphPad Software, version 8.4.2). All data were presented as mean ± standard error of the means (SEM). Statistical differences were assessed using two-way ANOVA test or two-tailed paired Student’s t-test.

### Single-cell RNA/TCR seq data processing

Single-cell expression was analyzed using the Cell Ranger single-cell Software Suite (v2.0.1 and v3.0.2 for the last 2 patients, 10X Genomics) to perform quality control, sample de-multiplexing, barcode processing, and single-cell 3’ and 5 ‘ gene counting. Sequencing reads were aligned to the GRCh38 human reference genome. We used emptyDrops (*39*) function from R package dropletUtils when analyzing the samples processed with version Cellranger v2.0.1. Cells with a p < 0.01 were considered for further analysis. The emptyDrops function is already implemented in Cellranger v3. Further analysis was performed in R (v3.5.1) using the Seurat package (v3.1.1) (*18*). Cells were then filtered out when expressing less than 500 genes or more than 5000 genes, or when expressing more than 10% mitochondrial genes. Altogether, among the 17 samples (11 from tumors, 6 from blood and 2 from juxta-tumor) 54247 cells were kept for statistical analysis. For each sample, the gene-cell-barcode matrix of the samples was then normalized to a total of 1e4 molecules. The top 2000 variable features were identified using the “vst” method from Seurat where both lowly and highly expressed genes are transformed onto a common scale. For the 11 tumor samples altogether, we then computed the integration anchors using the Seurat v3 integration method. This method is leveraging closely-related cells (termed anchors) between datasets to compute a batch-corrected matrix. Top 30 CCA components were used to find transfer anchors between datasets and used to generated the integrated matrix for the 11 tumors.

### Dimension reduction and unsupervised clustering

Top 30 Principal Components were computed and UMAP was performed using the top 30 PCs of the integrated matrix. Clusters were identified using the *FindNeighbors* and *FindClusters* function in Seurat with a resolution parameter of 1.2 and using the first 30 principal components. To choose the optimal number of clusters and prevent overclustering, clustree analysis was performed using the clustree package v0.4.1 (*40*). Unique cluster-specific genes were identified by running the Seurat *FindAllMarkers* function using Wilcoxon test on the uncorrected matrix. Then, three clusters containing contaminating cells were removed from the analysis: A cluster of myeloid cells expressing APOE, B cells expressing CD19, epithelial cells expressing KRT1. Signature scores were computed using the Seurat function *AddModuleScore* using the gene signature of interest and the integrated matrix. Briefly this function calculates for each individual cell the average expression of each gene signature, subtracted by the aggregated expression of control gene sets. The average heatmap was generated using the Average function in Seurat which is averaging gene expression across clusters. Cell cycle scoring was also performed using Seurat “CellCycleScore” function using cell cycle genes.

### Pathway enrichment

Pathways enrichment tests were performed using Metascape (*41*) with default parameters using differentially expressed genes between early precursors (CD8-KLF2, CD8-GZMK, CD8-XCL1) and terminally differentiated CD8^+^ T cells (CD8-GZMH, CD8-LAYN). Gene sets from MSigDB, v.7.0 were downloaded in GMT format from https://www.gsea-msigdb.org/gsea/msigdb/collections.jsp. These gene sets were used as modules for the “module score” function in Seurat.

### Label transfer using a reference

We chose to term “unassigned“ cells that had a prediction score below the bottom 5% of the positive control represented by the merging of 4 tumors in the 11 tumors reference.

Data from Guo et al. 2018 (*13*) was downloaded from GSE99254. Sade-Feldman et al. 2018 (*17*) data was downloaded from GSE120575. Azizi et al. 2018 (*16*) data was downloaded from GSE114727 and reprocessed using Seurat pipeline.

### Denoising using ENHANCE

We used ENHANCE (*42*), a denoising strategy based on principal component analysis (PCA) for visualization when genes were dropped-out (TOX and TCF7). This algorithm is base PCA to separate biological heterogeneity from technical noise and then mitigating the bias towards highly expressed genes by aggregating closely related cells. The algorithm was run with default parameters as described in the preprint. Since such denoising algorithms might lead to overcorrection, we still used the original matrix for the rest of the analysis.

### Silhouette score

Silhouette widths were computed using the first 30 PCs using the silhouette function from the R package cluster v2.1.0. The silhouette value is a measure of how similar a cell is to its own clusters compared to other clusters. The silhouette ranges from −1 to +1. A high value shows that the cell is well matched to its own cluster and poorly matched to neighboring clusters. A value around 0 indicates that the sample is close to the decision boundary between two neighboring clusters and negative values indicate that those samples might have been wrongly assigned to a cluster.

### Trajectory analysis

To compute pseudo-time alignment of our transcriptomes, we first used Monocle3 (v2.99.1) using the first 30 PCs of the integrated matrix to preform preprocessing and UMAP reduction. DDRTree algorithm was then used to reconstruct the tree embedding. PAGA output was generated using Scanpy v. 1.4.3 (*43*) with default values and a threshold of 0.15.

### TCR analysis

TCR-seq data for each sample was processed using Cell Ranger software (versions as above), with the command ‘cellranger vdj’ using the human reference genome GRCh38.

Cells with the same TCR receptor can be considered clonally related, however two factors complicate the identification of clonotypes:

- One cell can contain up to two alpha chains and one beta chain but they may not all be sequenced in a given cell due to e.g. dropout. This can lead to the artificial split of one clonotype into two or alternatively to the union of two different clonotypes which happen to share one chain by chance.
- Two unrelated cells can be barcoded together within the same droplet (a doublet), joining two independent clonotypes.

We choose a pragmatic, easily reproducible approach and use the same criteria as 10X: for all the samples collected from a patient, cells that share the exact same set of productive CDR3 nucleotide sequences are considered clones.

The STARTRAC expansion index, defined by Zheng et al. 2019 (*33*), it gives a measure of how much clonotypes in a given cluster have expanded. If a cluster *C* contains *N* clonotypes, the expansion index is given by:

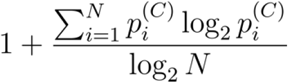

Where 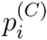 is the frequency of the cells of clonotype *i* in the cluster *C*.

Similarly, the STARTRAC transition index between two cluster *C*_1_ and *C*_2_, used in Fig 4F is defined by the formula:

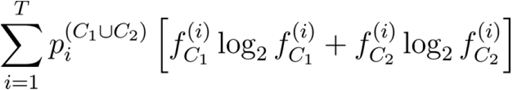

Where 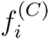 is the frequency of cells belonging to the cluster *C* inside the clonotype *i*.

The proximity measure between clusters that we have chosen to use in Fig 4D is given by:

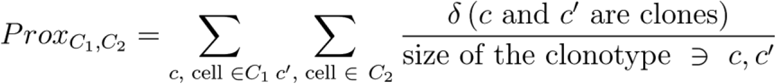

This measure is symmetric and it corresponds to the probability of finding a cell in *C*_2_ if one of its clones is in *C*_1_, normalized by the probability of the same thing happening when the clusters’ labels are randomly shuffled.

In every “circos” plot, the links between two clusters represent a clonotype shared between at least two different clusters. The start and end clusters correspond to the two most common clusters inside this clonotype.

## Supplementary Materials

Fig. S1. Comparison of the 3’ vs. 5’ single cell chemistries and general characterization of total CD3^+^ T cell subsets.

Fig. S2. Gene signatures and biological pathways differentiating precursors and terminally differentiated CD8^+^ T cells.

Fig. S3. Flow cytometry gating strategy and quantification of silhouette score and maximum prediction score of the label transfer method in CD8^+^ T cell subsets.

Fig. S4. Quantification of shared clones among the CD8^+^ clusters and transition index.

Fig. S5. Characterization of the cycling CD8^+^ T cells.

Fig. S6. Analysis of tissue-specificity of tumor, juxta and blood tissue in NSCLC patients.

Table S1. Differentially expressed genes per cluster.

Table S2. Gene signatures.

## Acknowledgments

The authors wish to thank the Flow Cytometry Platform of Institut Curie and Lea Guyonnet for helpful discussions. We thank all the thoracic surgery, respiratory medicine, pathology, and tissue procurement staff of Institut Mutualiste Montsouris, as well as all the patients and their families.

## Funding

C.M received a PhD fellowship from Servier Laboratories and funds by the PhD Program “FIRE - Programme Bettencourt”. P.G. is supported by a PhD fellowship from Ligue Nationale contre le Cancer at Université Paris Descartes (Paris V). This study was supported by grants from “Janssen Horizon”, “Fondation ARC pour la recherche sur le cancer”, “Fondation chercher et trouver - StiftungSchweiz”, “European Research Council COG 724208” (T.D, T.M & A.W), ANR-10-EQPX-03 (Equipex), ANR-10-INBS-09-08 (France Génomique Consortium) from the Agence Nationale de la Recherche (“Investissements d’Avenir” program) and the SiRIC-Curie program - SiRIC Grant INCa-DGOS-4654.

## Author contributions

S.A, J.W and O.L conceived the project. C.M and M.L designed & performed experiments and acquired the data. P.G and C.M analyzed and interpreted the data. T.D analyzed the data. T.M, A.W, C.G, J.W and S.A guided computational analysis. S.B and S.L were involved in data acquisition. N.G, A.S and M.L were involved in patient inclusion and sample acquisition. C.M, P.G, J.W and S.A wrote the manuscript with contributions from all authors. J.W and S.A supervised the project.

## Competing interests

The authors declare that they have no competing interests.

## Notes

### Competing Interest Statement

The authors have declared no competing interest.

