## Supplementary figures for "Contribution of resident and circulating precursors to tumor-infiltrating CD8^+^ T cell populations in non-small cell lung cancer patients"

**Supplementary Data**

**Figure S1. Comparison of the 3' vs. 5' single cell chemistries and general characterization of total CD3$^{+}$ T cell subsets. (A)** UMAP representation of the 11 samples without integration. The strongest batch component can be explained by the 3' vs. 5' -oriented single cell chemistries. **(B)** PCA representation of the bulk-like 11 samples for the original or corrected matrices. **(C)** Quantification of variance explained by PCs for the uncorrected and batch-corrected matrices. Batch-correction is able to reduce the impact of single-cell chemistries. **(D)** Projection of single gene markers, identifying T cell populations. Each cell is colored based on the normalized gene expression. **(E)** UMAP projection of cell density onto the CD3$^{+}$ T cell dataset from the 11 NSCLC tumor samples showing continuous states of differentiation. **(F)** Heatmap of averaged expression values for the inhibitory and stimulating immune checkpoint markers in all T cell clusters. **(G)** Heatmap of cell fractions distribution per cluster in each patient.

**Figure S2. Gene signatures and biological pathways differentiating precursors and terminally differentiated CD8$^{+}$ T cells. (A)** UMAP projection of tumor infiltrating T cells, with each cell colored based on the relative normalized expression of a gene signature[19] for exhausted progenitors (left panel) or terminal exhausted cells (right panel). **(B)** Pathway analysis for precursor CD8$^{+}$ T cells (left panel) and terminally differentiated CD8$^{+}$ cells (right panel). Analysis was performed based on the top DEGs for each of the populations. **(C)** UMAP and quantification of clusters after label transfer of CD3$^{+}$ T cells from 3 different tumor types (NSCLC[13], Breast[16], Melanoma[17]) label transferred onto our 11 CD3$^{+}$ single-cell references

**Figure S3. Flow cytometry gating strategy and quantification of silhouette score and maximum prediction score of the label transfer method in CD8$^{+}$ T cell subsets. (A)** UMAP projection of tumor infiltrating T cells. Each cell is colored based on the normalized gene expression of KLRG1 (left panel) or ZNF683 (right panel). **(B)** Gating strategy for CD8$^+$ tumor-infiltrating cells. CD8$^+$ cells are divided in 8 subpopulations based on the expression of PD-1, TIM3 and CD39: PD-1$^-$, TIM3$^-$, CD39$^-$ triple-negative (TN), PD-1$^+$ (TIM3$^-$ CD39$^-$), TIM3$^+$ (PD-1$^-$ CD39$^-$), CD39$^+$ (PD-1$^-$TIM3$^-$), TIM3$^+$ CD39$^+$ (PD-1$^-$), PD-1$^+$ TIM3$^+$ CD39$^+$ -triple-positive (TP). **(C)** Clustree information overlayed onto the UMAP embedding. The early separation (res 0.2) between the resident and circulating part of CD8$^+$ indicates that these 2 subparts of the datasets are the strongest difference within the data. **(D)** Feature plot of marker genes overlayed onto the Monocle pseudo-time model. We recapitulate the results from the original UMAP embedding. **(E)** Distribution of silhouette score in CD8$^+$ T cell clusters. **(F)** Distribution of maximum prediction score of the label transfer method for CD8-GZMH cluster into the rest CD8$^+$ T cell clusters

**Figure S4. Quantification of shared clones among the CD8$^+$ clusters and transition index. (A)** Distribution of top clonotypes per patients and colored by clusters identities. Each line corresponds to a clone. **(B)** Transition index (See methods). **(C)** Quantification of the number of shared clones among the CD8+ clusters **(D)** Scatter plots of clonal sharing among CD8+ clusters. Each dot corresponds to a clone.

**Figure S5. Characterization of the cycling CD8$^+$ T cells. (A)** Violin plots for G2M (top) and S phase (bottom) signature score according to CD3+ clusters. The 2 cycling clusters are split by the cycling phase. **(B)** tSNE visualization of additional markers, showing Ki67 marker co-expression with CD103$^+$CD39$^+$TIM3$^+$ CD8$^+$ subset. **(C)** Volcano plot of cycling GZMH versus cycling LAYN. DNA repair genes are being upregulated in the cycling LAYN part of the terminally differentiated cells. **(D)** Violin plots showing CD8 LAYN signature from Guo et al.[13] in cycling vs non cycling (cycling / non cycling computed using label transfer). **(E)** Feature plots of corresponding LAYN and cycling signature in Guo et al. [13] dataset. **(F)** Scatter plots representing

TCR clones cluster identities. Number of cycling cells according to the number of cells in the other CD8 subsets. The size of each circle is proportional to the clonal size of each clone. **(G)** Ternary plot showing relationships between percentage of cycling cells, CD8-GZMH and its neighboring clusters.

**Figure S6. Analysis of tissue-specificity of tumor, juxta and blood tissue in NSCLC patients.** **(A)** Unsupervised clustering of the 3 integrated tissues: tumor, juxta-tumor tissue and blood. **(B)** Frequency of clusters split by tissue and by patient for every cluster. **(C)** Flow cytometry data showing the frequency of CD8+ T cells subsets in 3 different tissues in a dataset of 5 NSCLC patients (5 matched tumor and blood samples, 2 matched tumor, juxta-tumor and blood samples).

**Fig. S1.**

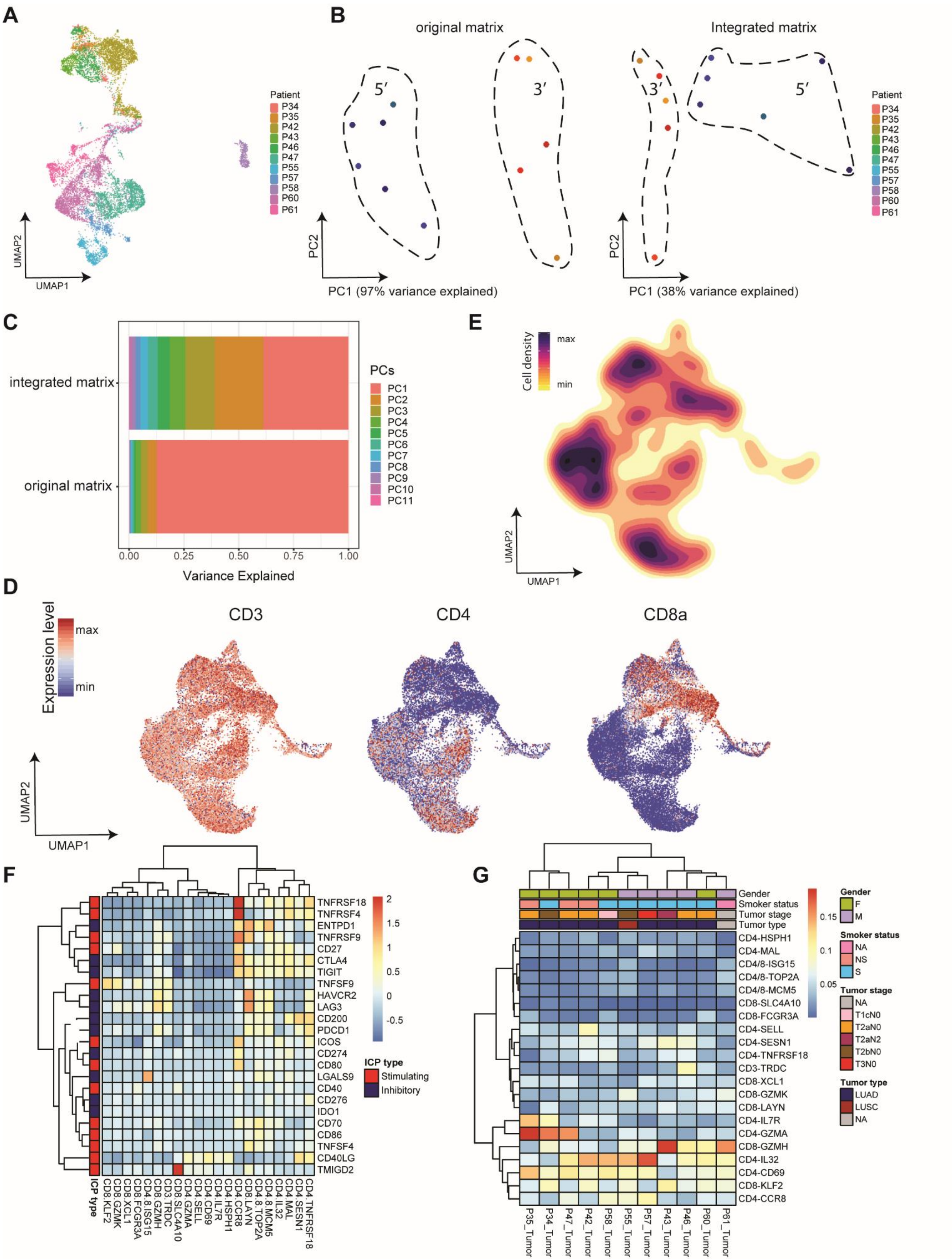
A
B
original matrix
Integrated matrix
5′
3′
Patient
P34
P35
P42
P43
P46
P47
P55
P57
P58
P60
P61
UMAP2
UMAP1
PC2
PC1 (97% variance explained)
PC1 (38% variance explained)
C
integrated matrix
original matrix
Variance Explained
PCs
PC1
PC2
PC3
PC4
PC5
PC6
PC7
PC8
PC9
PC10
PC11
E
Cell density
max
min
D
Expression level
CD3
CD4
CD8a
F
ICP type
Stimulating
Inhibitory
G
Gender
Smoker status
Tumor stage
Tumor type
LUAD
LUSC
NA

**Fig. S2.**

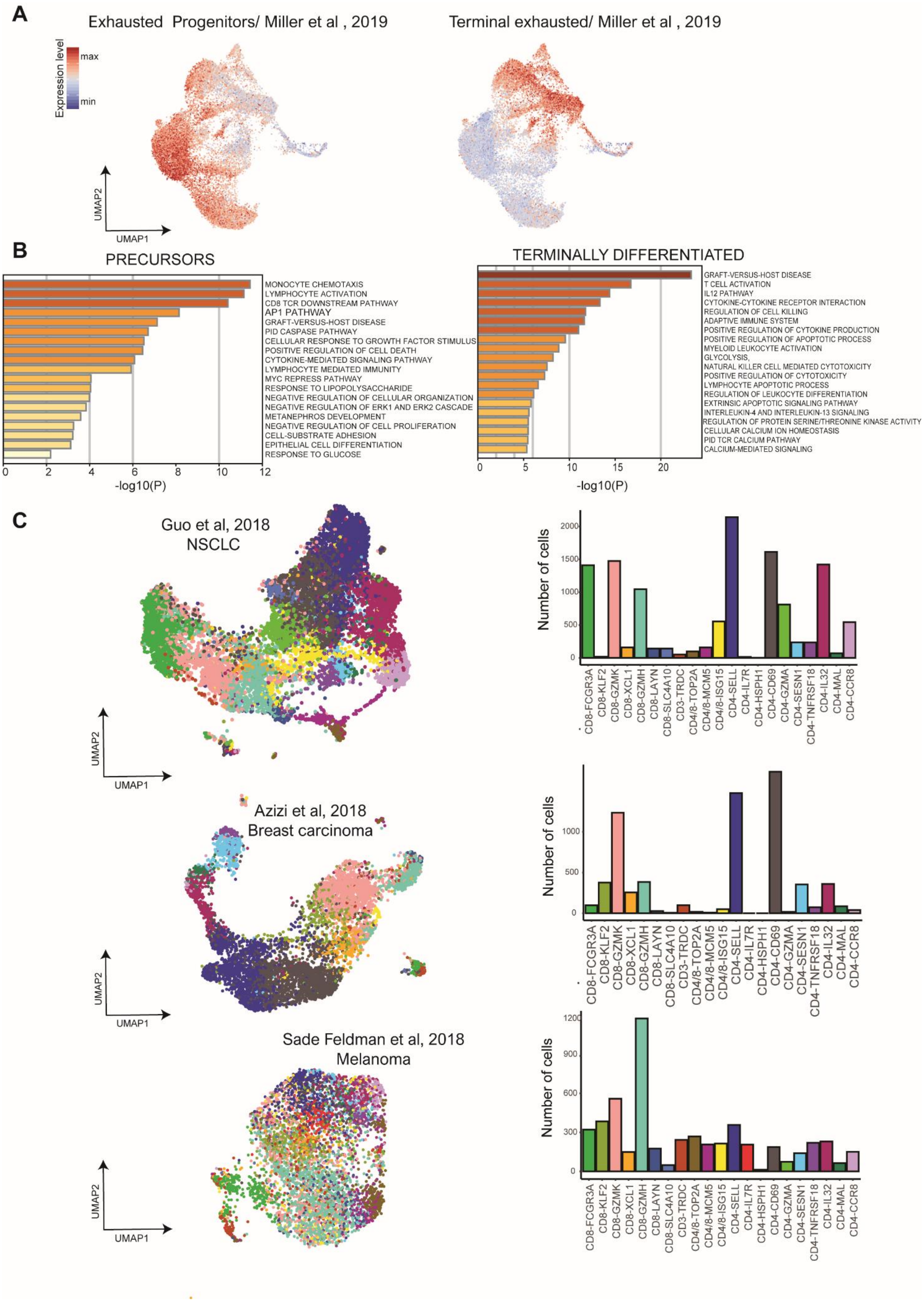

A
Exhausted Progenitors/ Miller et al , 2019
Terminal exhausted/ Miller et al , 2019
Expression level
max
min
UMAP2
UMAP1
B
PRECURSORS
MONOCYTE CHEMOTAXIS
LYMPHOCYTE ACTIVATION
CD8 TCR DOWNSTREAM PATHWAY
AP1 PATHWAY
GRAFT-VERSUS-HOST DISEASE
PID CASPASE PATHWAY
CELLULAR RESPONSE TO GROWTH FACTOR STIMULUS
POSITIVE REGULATION OF CELL DEATH
CYTOKINE-MEDIATED SIGNALING PATHWAY
LYMPHOCYTE MEDIATED IMMUNITY
MYC REPRESS PATHWAY
RESPONSE TO LIPOPOLYSACCHARIDE
NEGATIVE REGULATION OF CELLULAR ORGANIZATION
NEGATIVE REGULATION OF ERK1 AND ERK2 CASCADE
METANEPHROS DEVELOPMENT
NEGATIVE REGULATION OF CELL PROLIFERATION
CELL-SUBSTRATE ADHESION
EPITHELIAL CELL DIFFERENTIATION
RESPONSE TO GLUCOSE
0 2 4 6 8 10 12
-log10(P)
TERMINALLY DIFFERENTIATED
GRAFT-VERSUS-HOST DISEASE
T CELL ACTIVATION
IL12 PATHWAY
CYTOKINE-CYTOKINE RECEPTOR INTERACTION
REGULATION OF CELL KILLING
ADAPTIVE IMMUNE SYSTEM
POSITIVE REGULATION OF CYTOKINE PRODUCTION
POSITIVE REGULATION OF APOPTOTIC PROCESS
MYELOID LEUKOCYTE ACTIVATION
GLYCOLYSIS,
NATURAL KILLER CELL MEDIATED CYTOTOXICITY
POSITIVE REGULATION OF CYTOTOXICITY
LYMPHOCYTE APOPTOTIC PROCESS
REGULATION OF LEUKOCYTE DIFFERENTIATION
EXTRINSIC APOPTOTIC SIGNALING PATHWAY
INTERLEUKIN-4 AND INTERLEUKIN-13 SIGNALING
REGULATION OF PROTEIN SERINE/THREONINE KINASE ACTIVITY
CELLULAR CALCIUM ION HOMEOSTASIS
PID TCR CALCIUM PATHWAY
CALCIUM-MEDIATED SIGNALING
0 5 10 15 20
-log10(P)
C
Guo et al, 2018
NSCLC
Azizi et al, 2018
Breast carcinoma
Sade Feldman et al, 2018
Melanoma
Number of cells
CD8-FCGR3A
CD8-KLF2
CD8-GZMK
CD8-XCL1
CD8-GZMH
CD8-LAYN
CD8-SLC4A10
CD3-TRDC
CD4/8-TOP2A
CD4/8-MCM5
CD4/8-ISG15
CD4-SELL
CD4-IL7R
CD4-HSPH1
CD4-CD69
CD4-GZMA
CD4-SESN1
CD4-TNFRSF18
CD4-IL32
CD4-MAL
CD4-CCR8

**Fig. S3.**

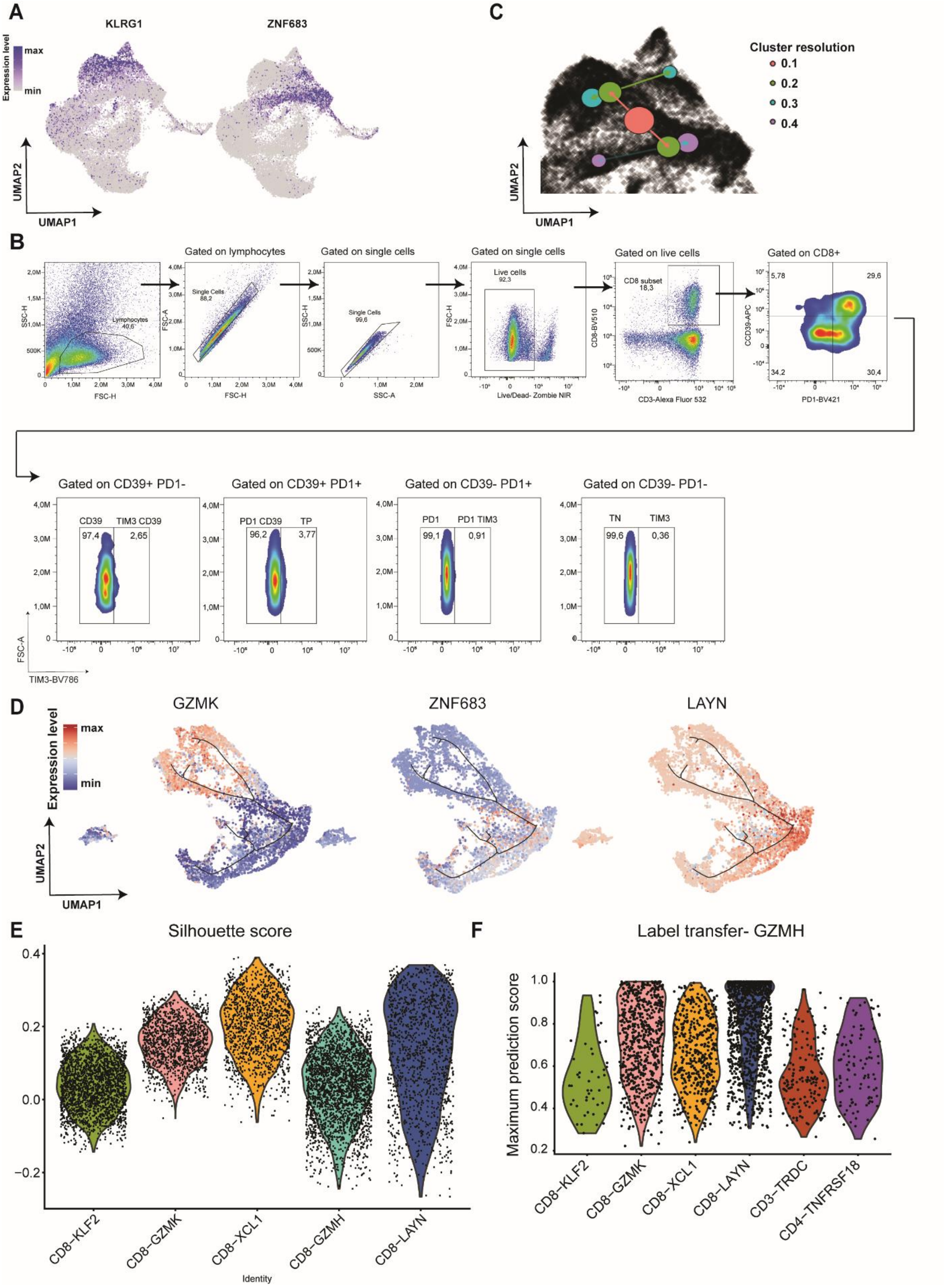

A
KLRG1
ZNF683
Expression level
max
min
UMAP2
UMAP1
C
Cluster resolution
0.1
0.2
0.3
0.4
B
Gated on lymphocytes
Gated on single cells
Gated on single cells
Gated on live cells
Gated on CD8+
Lymphocytes 40,6
Single Cells 88,2
Single Cells 99,6
Live cells 92,3
CD8 subset 18,3
FSC-H
SSC-H
FSC-A
SSC-A
Live/Dead- Zombie NIR
CD8-BV510
CD3-Alexa Fluor 532
CD39-APC
PD1-BV421
5,78
29,6
34,2
30,4
Gated on CD39+ PD1-
Gated on CD39+ PD1+
Gated on CD39- PD1+
Gated on CD39- PD1-
CD39
TIM3 CD39
97,4
2,65
PD1 CD39
TP
96,2
3,77
PD1
PD1 TIM3
99,1
0,91
TN
TIM3
99,6
0,36
FSC-A
TIM3-BV786
D
GZMK
ZNF683
LAYN
Expression level
max
min
UMAP2
UMAP1
E
Silhouette score
CD8−KLF2
CD8−GZMK
CD8−XCL1
CD8−GZMH
CD8−LAYN
Identity
F
Label transfer- GZMH
Maximum prediction score
CD8−KLF2
CD8−GZMK
CD8−XCL1
CD8−LAYN
CD3−TRDC
CD4−TNFRSF18

Fig. S4.

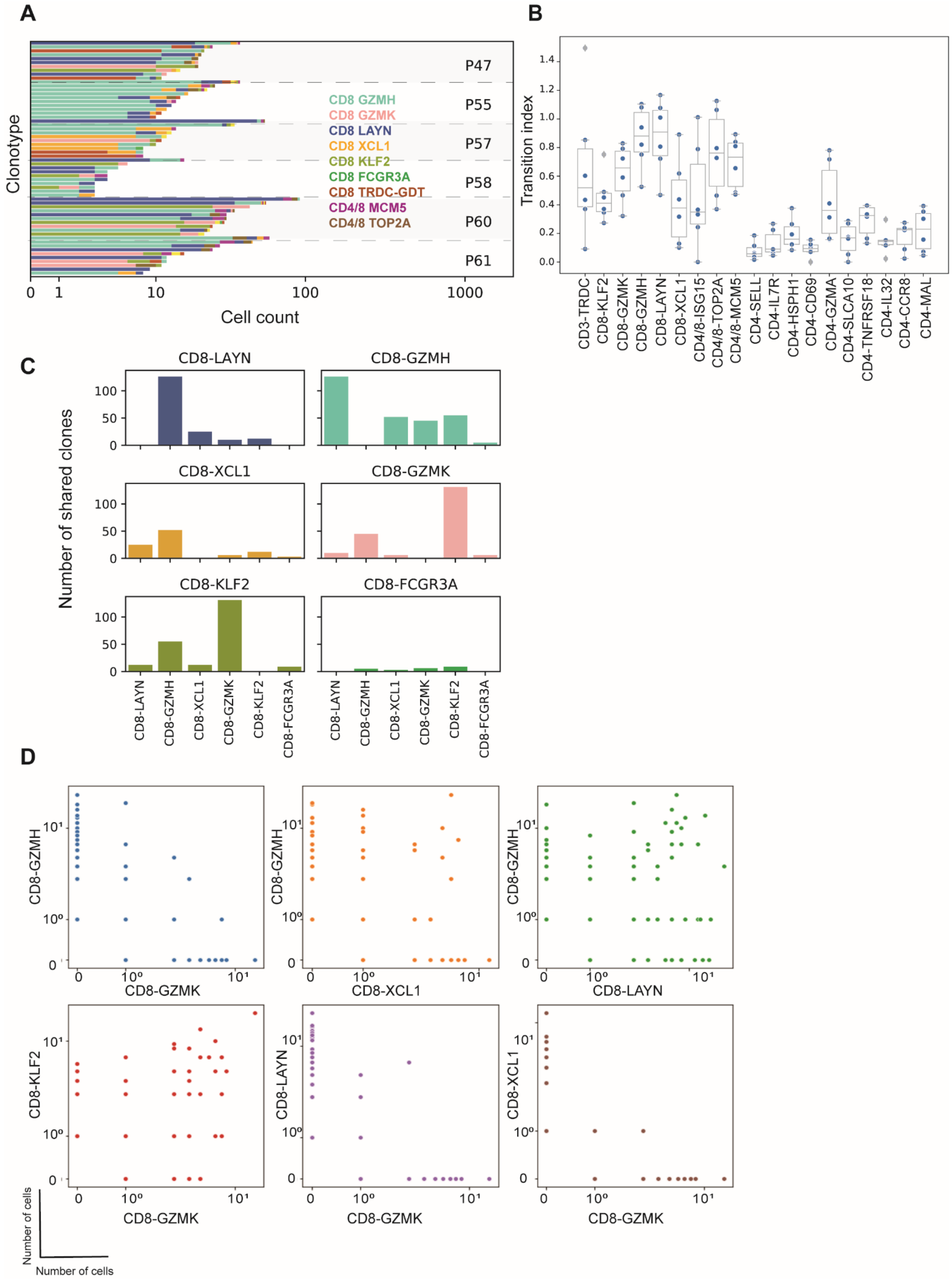
A
B
C
D
Clonotype
Cell count
P47
P55
P57
P58
P60
P61
CD8 GZMH
CD8 GZMK
CD8 LAYN
CD8 XCL1
CD8 KLF2
CD8 FCGR3A
CD8 TRDC-GDT
CD4/8 MCM5
CD4/8 TOP2A
Transition index
CD3-TRDC
CD8-KLF2
CD8-GZMK
CD8-GZMH
CD8-LAYN
CD8-XCL1
CD4/8-ISG15
CD4/8-TOP2A
CD4/8-MCM5
CD4-SELL
CD4-IL7R
CD4-HSPH1
CD4-CD69
CD4-GZMA
CD4-SLCA10
CD4-TNFRSF18
CD4-IL32
CD4-CCR8
CD4-MAL
Number of shared clones
CD8-LAYN
CD8-GZMH
CD8-XCL1
CD8-GZMK
CD8-KLF2
CD8-FCGR3A
CD8-GZMK
CD8-XCL1
CD8-LAYN
CD8-GZMH
CD8-KLF2
Number of cells
Number of cells

**Fig. S5**

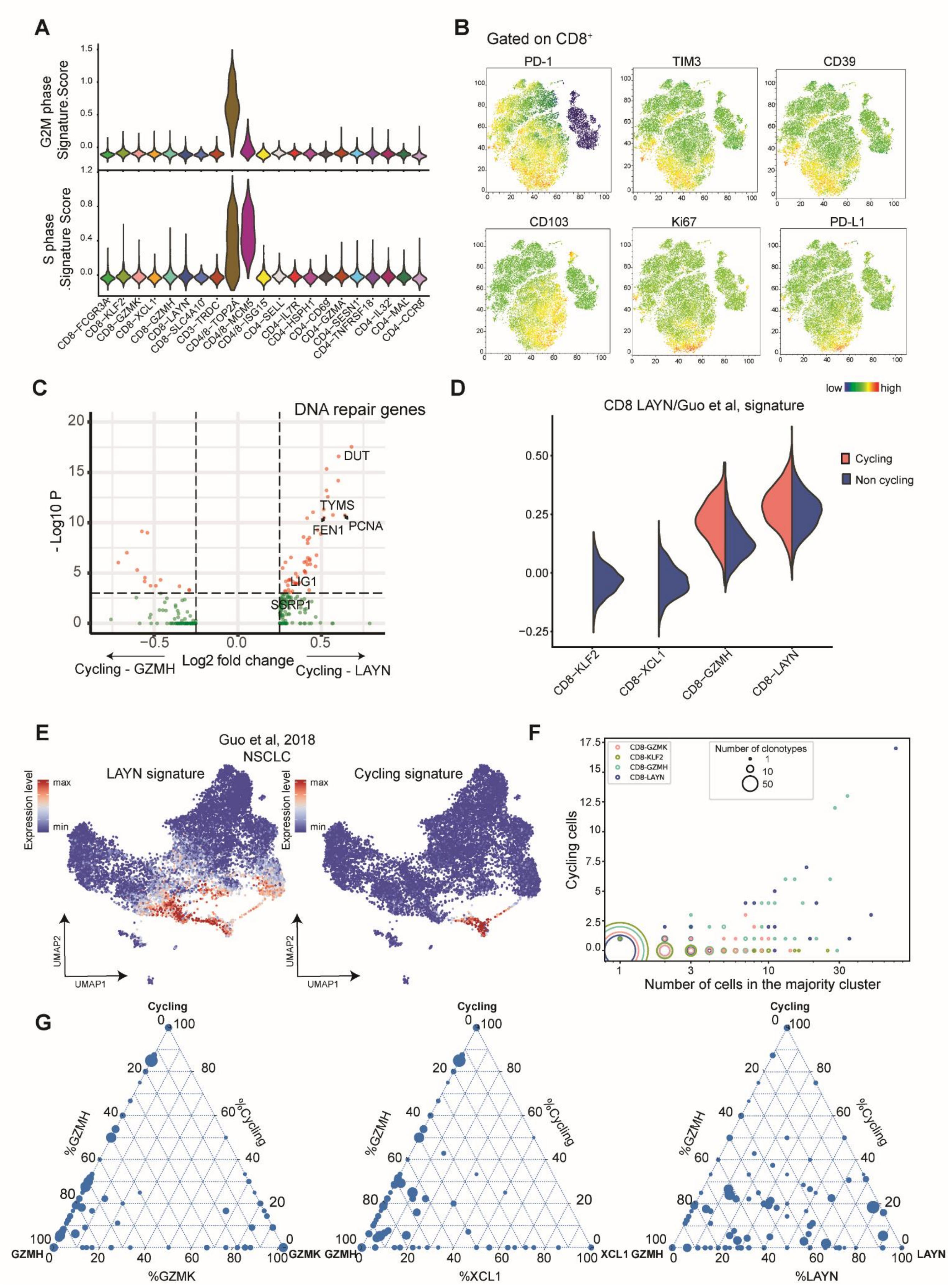
A
G2M phase Signature.Score
S phase Signature Score
CD8−FCGR3A
CD8−KLF2
CD8−GZMK
CD8−XCL1
CD8−GZMH
CD8−LAYN
CD8−SLC4A10
CD3−TRDC
CD4/8−TOP2A
CD4/8−MCM5
CD4/8−ISG15
CD4−SELL
CD4−IL7R
CD4−HSPH1
CD4−CD69
CD4−GZMA
CD4−SESN1
CD4−TNFRSF18
CD4−IL32
CD4−MAL
CD4−CCR8
B
Gated on CD8+
PD-1
TIM3
CD39
CD103
Ki67
PD-L1
low
high
C
DNA repair genes
- Log10 P
Log2 fold change
DUT
TYMS
FEN1
PCNA
LIG1
SSRP1
Cycling - GZMH
Cycling - LAYN
D
CD8 LAYN/Guo et al, signature
Cycling
Non cycling
CD8−KLF2
CD8−XCL1
CD8−GZMH
CD8−LAYN
E
Guo et al, 2018
NSCLC
LAYN signature
Cycling signature
Expression level
max
min
UMAP1
UMAP2
F
CD8-GZMK
CD8-KLF2
CD8-GZMH
CD8-LAYN
Number of clonotypes
Cycling cells
Number of cells in the majority cluster
G
Cycling
%GZMH
%Cycling
GZMH
GZMK
%GZMK
XCL1
%XCL1
LAYN
%LAYN

**Fig. S6.**

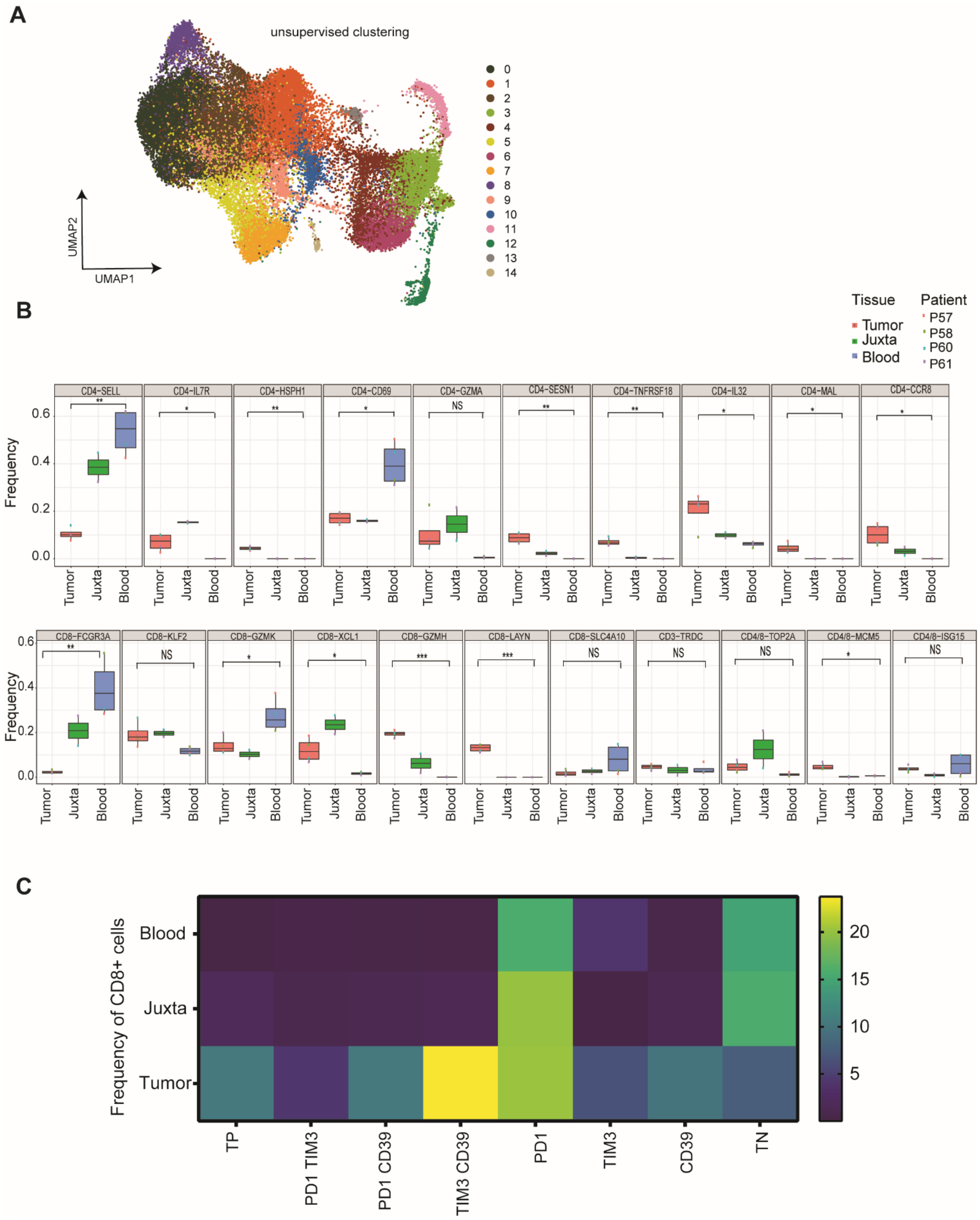
A
unsupervised clustering
UMAP2
UMAP1
B
Tissue
Tumor
Juxta
Blood
Patient
P57
P58
P60
P61
Frequency
CD4-SELL
CD4-IL7R
CD4-HSPH1
CD4-CD69
CD4-GZMA
CD4-SESN1
CD4-TNFRSF18
CD4-IL32
CD4-MAL
CD4-CCR8
CD8-FCGR3A
CD8-KLF2
CD8-GZMK
CD8-XCL1
CD8-GZMH
CD8-LAYN
CD8-SLC4A10
CD3-TRDC
CD4/8-TOP2A
CD4/8-MCM5
CD4/8-ISG15
NS
C
Frequency of CD8+ cells
Blood
Juxta
Tumor
TP
PD1 TIM3
PD1 CD39
TIM3 CD39
PD1
TIM3
CD39
TN
20
15
10
5

**Table S1. Differentially expressed genes per cluster.** List of DEGs per cluster with average log fold change information.

**Table S2. Gene signatures.** List of all gene signatures having been used in the data analysis.
